# Accurate and efficient constrained molecular dynamics of polymers through Newton’s method and special purpose code

**DOI:** 10.1101/2022.09.28.509839

**Authors:** Lorién López-Villellas, Carl Christian Kjelgaard Mikkelsen, Juan José Galano-Frutos, Santiago Marco-Sola, Jesús Alastruey-Benedé, Pablo Ibáñez, Miquel Moretó, Javier Sancho, Pablo García-Risueño

## Abstract

In molecular dynamics simulations we can often increase the time step by imposing constraints on internal degrees of freedom, such as bond lengths and bond angles. This allows us to extend the length of the time interval and therefore the range of physical phenomena that we can afford to simulate. In this article we analyse the impact of the accuracy of the constraint solver. We present ILVES-PC, an algorithm for imposing constraints on proteins accurately and efficiently.

ILVES-PC solves the same system of differential algebraic equations as the celebrated SHAKE algorithm, but uses Newton’s method for solving the nonlinear constraint equations. It solves the necessary linear systems of equations using a specialised linear solver that utilises the molecular structure. ILVES-PC can rapidly solve the nonlinear constraint equations to nearly the limit of machine precision. This eliminates the spurious forces introduced to simulations through the very common use of inaccurate approximations. The run-time of ILVES-PC is proportional to the number of constraints.

We have integrated ILVES-PC into GROMACS and simulated proteins of different sizes. Compared with SHAKE, we have achieved speedups of up to 4.9× in single-threaded executions and up to 76× in shared-memory multi-threaded executions. Moreover, we find that ILVES-PC is more accurate than the P-LINCS algorithm. Our work is a proof-of-concept of the utility of software designed specifically for the simulation of polymers.

**Author summary:** Molecular dynamics simulates the time evolution of molecular systems. It has become a tool of extraordinary importance for e.g. understanding biological processes and designing drugs and catalysts. This article presents an algorithm for computing the forces needed to impose constraints in molecular dynamics, i.e., the *constraint forces*; moreover, it analyses the effect of the accuracy of the constraint solver. Presently, it is customary to calculate the constraint forces with a relative error that that is not tiny. This is due to the high computational cost associated with the available software. Accurate calculations are possible, but they are very time-consuming. The algorithm that we present solves this problem: it computes the constraint forces accurately and efficiently. Our work will improve the accuracy and reliability of molecular dynamics simulations beyond the present state-of-the-art. The results that we present are also a proof-of-concept that special-purpose code can increase the performance of software for the simulation of polymers. The algorithm is implemented into a popular molecular simulation package, and is now available for the research community.

## 1 Introduction

Molecular simulation is a powerful research tool for scientific and technological purposes. It is applied to a wide range of problems in chemistry and biology, such as the development of novel materials [1] or biomedicines, e.g., for fighting cancer [2] and infectious diseases, like the SARS-CoV-2 [3, 4]. One of the most widely used techniques for molecular simulations is molecular dynamics (MD) [5, 6], which calculates the time evolution of molecular systems subject to the Newton’s equations, thus enabling the calculation of a variety of quantities whose measurement in laboratories is frequently either difficult or unfeasible. The impact of molecular simulation is expected to increase greatly due to the continuous improvement of available computational capabilities [7] and calculation methods [8]. Among the former, we highlight the successive generations of the Anton supercomputers [9]; among the latter, the solution of the protein folding problem by AlphaFold [10]. The availability of 3D structures of proteins provided by AlphaFold will probably boost their simulations, e.g., for analysing their capabilities as catalysts or medicines or for a more accurate interpretation of the effect of mutations on the phenotype [11]. The availability of accurate and efficient methods for such simulations does hereby acquire a novel boost.

## 2 Background and Motivation

### 2.1 Notation

All vectors are written using bold lowercase letters. All vectors are column vectors by default. When we need a row vector, then we shall explicitly transpose a column vector. The Euclidean norm of a vector ***x*** = (*x*_1_, *x*_2_,…, *x*_*n*_)^*T*^ *∈* ℝ_*n*_ is written as ‖***x****‖* and is the nonnegative number ‖*x*‖ given by 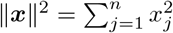. All matrices are written using bold uppercase letters. If the function ***f*** = (*f*_1_, *f*_2_,…, *f*_*n*_)^*T*^ : ℝ_*n*_ *→* ℝ_*n*_ is differentiable, then the Jacobian ***F*** (***x***) of ***f*** at the point ***x*** *∈* ℝ_*n*_ is the matrix ***A*** = [*a*_*ij*_] *∈* ℝ^*n*×*n*^ given by

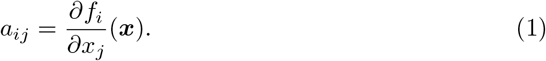

We shall now define the notation used to describe a system of *m* atoms. Let *m*_*i*_ *>* 0 denote mass of the *i*th atom and let ***m***_*i*_ *∈* ℝ^3^ and the diagonal mass matrix ***M*** *∈*ℝ^3*m*×3*m*^ be given by

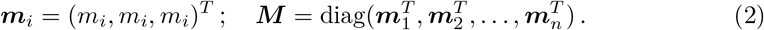

In addition, let ***q***_*i*_, ***v***_*i*_, ***f***_*i*_ *∈* ℝ^3^ denote the position of, the velocity of, and the force acting on the *i*th atom, and let ***q*** *∈* ℝ^3*m*^, ***v*** *∈* ℝ^3*m*^, and ***f*** *∈* ℝ^3*m*^ be given by

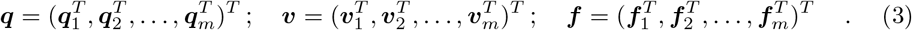

### 2.2 Fundamentals of constrained molecular dynamics

By Newton’s second law, the equations of motion of a system of *m* atoms are

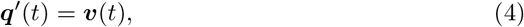

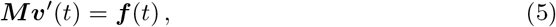

where the prime indicates differentiation with respect to the time *t*. In nature, the motion of atoms is continuous in time; however a computer simulation customarily uses a sequence of discrete time steps. The standard algorithm for this problem is the velocity Verlet-algorithm [12]. It is well-known that certain motions such as bond stretching, bond bending, and torsional vibrations are all periodic with characteristic frequencies that depend on the atoms involved [13]. It is generally accepted that in order to accurately resolve periodic motion one needs at least five time steps per period. Hence the fastest vibration imposes a limitation on the maximum time step that can be used and this limits the length of the time interval one can afford to simulate. In order to simulate phenomena with a longer duration it is customary to constrain the fastest degrees of freedom. Let *n* denote the number of constraints. Mathematically, the problem consists of solving the following system of differential algebraic equations

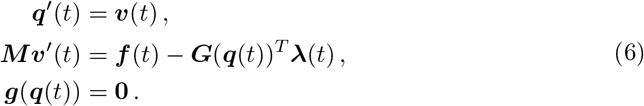

with respect to ***q, v***, and ***λ***. Here ***g*** : ℝ^3*m*^ *→* ℝ^*n*^ is the constraint function, i.e.,

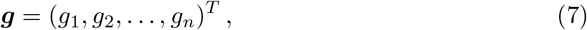

where *g*_*i*_ : ℝ^3*m*^ *→* ℝ is the *i*th constraint function and ***G***(***q***) *∈* ℝ^*n*×3*m*^ is the Jacobian of ***g*** at the point ***q***. The vector −***G***(***q***(*t*))^*T*^ ***λ***(*t*) is the *constraint force*.

### 2.3 Constrained MD solvers

Numerous algorithms for constrained molecular dynamics have been proposed [14–23]. Their main objective has been the reduction of the time-to-solution of the constraint equations. The most popular algorithms are SHAKE [24] and (P-)LINCS [25]. The SHAKE algorithm solves the system of differential algebraic equations (6) using a pair of staggered uniform grids with *fixed* time step *h>* 0. SHAKE’s equations take the form:

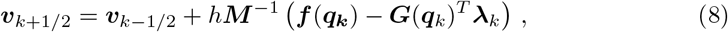

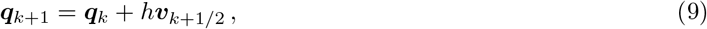

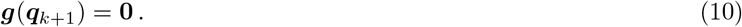

Here ***q***_*k*_ ≈ ***q***(*t*_*k*_) and 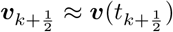, where *t*_*k*_ = *kh* and 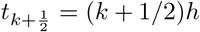. Equation (10) is a nonlinear equation for the unknown Lagrange multiplier ***λ***_*k*_, namely

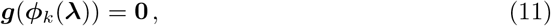

where ***ϕ***_*k*_ is the function given by

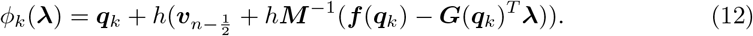

It is known that SHAKE is second order accurate in the time step [12]. The original SHAKE algorithm solved the constraint equations using the nonlinear Gauss-Seidel method, which converges locally and linearly subject to certain mild conditions (see [26] and the references therein). The LINCS and P-LINCS algorithms use a truncated Neumann-series to approximate the solution of the relevant linear systems. The Neumann-series converges linearly at best and there are physically relevant cases for which it does not converge at all [25, 27, 28]. Therefore, solving the constraints to the limit of machine precision is time-consuming. Indeed, as we shall demonstrate, the time spent by SHAKE can easily exceed 50% of the total execution time in realistic simulations; other research works also indicate severe performance drops (23%) when LINCS is accurately solved [78]. Thus, it would be useful to have software that solves the constraint equations accurately and rapidly. To this end, we have developed and implemented an algorithm called *ILVES* [29, 30] that avoids coarse-grain approximations and calculates the constraint forces accurately. We expect that avoiding coarse approximations will also produce a solver that is capable of finding solutions when the atomic displacements are more abrupt, e.g., during simulations run at high temperatures or with large time steps (like those used in Brownian dynamics [31] calculations), or when constraints on bond angles are also imposed. For instance, in Ref. [28] it was shown that 7200 distinct simulations all produced matrices for which the expansion used by LINCS does not converge and SHAKE had to be used instead. However, the SHAKE algorithm is commonly described as inherently serial [32], which makes it inefficient for the –presently ubiquitous– parallel computations. Since the parallel algorithm ILVES-PC solves the exact same equations as the sequential algorithm SHAKE, we expect that ILVES-PC will be able to solve more problems than LINCS. To sum up, ILVES combines the features of efficiency, parallelism, accuracy, and reliability.

We emphasize that several authors have already applied Newton’s method for solving nonlinear equations in the context of constrained molecular dynamics. M-SHAKE [20] treats the linear systems as dense and solves them using Gaussian elimination. This approach is limited to small molecules because the time complexity for computing an LU factorization of a dense matrix of dimension *n* is *O*(*n*^3^). MILC-SHAKE [17] utilizes the linear structure of alkanes to achieve a time complexity of *O*(*n*) by computing an LU factorization of a tridiagonal matrix rather than a fully dense matrix. The authors of the papers [21, 33] all approximate the relevant matrices using a matrices that are symmetric positive semi-definite and apply the conjugate gradient (CG) algorithm to solve these systems. The main advantages of this approach are twofold: the simplicity of the parallelisation of the CG algorithm, and the solution can be accelerated using a preconditioner. The disadvantage of this approach is the difficulty of finding a preconditioner whose quality can be guaranteed mathematically.

We shall now describe how Newton’s method can be applied in the context of molecular dynamics with constraints. We begin by stating the method in the case of a general nonlinear equation. Let ***f*** : ℝ^*n*^ *→* ℝ^*n*^ be a differential function and consider the problem of solving

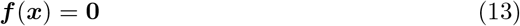

with respect to ***x*** *∈* ℝ^*n*^. If the Jacobian ***F*** of ***f*** is nonsingular, then Newton’s method is defined and takes the form

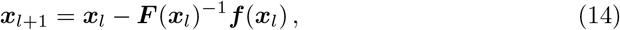

where the initial value ***x***_0_ must be chosen by the user. In general, we expect that Newton’s method will converge locally to a zero of ***f*** and that the convergence will be quadratic. In practice, we should never explicitly invert the matrix ***F*** (***x***_*l*_), instead we should compute the correction ***F*** (***x***_*l*_)^−1^***f*** (***x***_*l*_) by *solving* the linear system

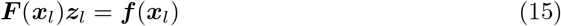

with respect to ***z***_*l*_. We emphasize this point by restating Newton’s method as the iteration

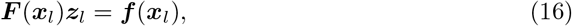

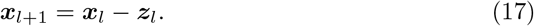

We now return to the nonlinear constraint equation (10). To this end, we introduce the matrix function ***A*** : ℝ^3*m*^ × ℝ^3*m*^ *→* ℝ^*n*×*n*^ given by

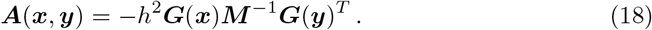

Then Newton’s method for the Lagrange multiplier ***λ***_*k*_ is given by

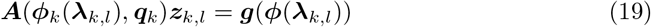

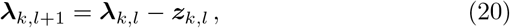

where the initial value ***λ***_*k*,0_ must be chosen by the user. The simple choice of ***λ***_*k*,0_ = **0** is the de-facto standard choice.

We mention in passing that the matrix ***A***(***x, y***) is structurally symmetric and that the matrix ***A***(***ϕ***_*k*_(***λ***_*k,l*_), ***q***_*k*_) is close to the symmetric matrix ***A***(***q***_*k*_, ***q***_*k*_), simply because

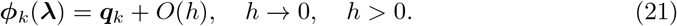

This is the observation that was utilized by the authors of the papers [21, 33].

### 2.4 Bond constraints

We now limit the discussion to general bond-length constraints. Note that constraints on bond angles are commonly enforced by constraining distances between two atoms. ILVES’ design is expected to provide accurate solutions when imposing any kind of constraints, either on bond lengths, bond angles or dihedral angles, due to the fact that the coordinate matrix ***A*** is banded (regardless of the kind of the constrained degrees of freedom) in biological molecules. Constraining degrees of freedom other than bond lengths using flexible constraints [34] may be an appropriate method to increase the time step of the simulation.

Our objective is to present a formula for the entries of the matrix ***A***(***x, y***). Let the *i*th bond have length *σ*_*i*_ *>* 0 and let *a*_*i*_ and *b*_*i*_ denote the indices of the two bonded atoms. Then the *i*th constraint can be written as

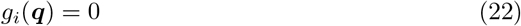

where

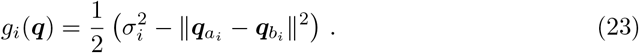

A direct calculation establishes that

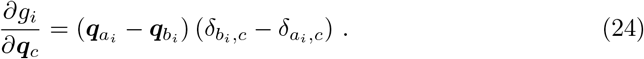

Here *δ*_*i*_ is Kronecker’s delta, i.e,

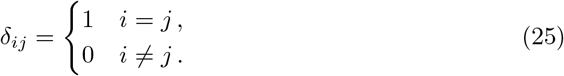

In particular, we observe that

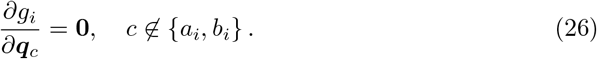

Now let *i* and *j* denote the indices of two bonds. There are exactly 3 distinct possibilities:

1. The two bonds have no atoms in common.
2. The two bonds share a single atom.
3. The two bonds are identical, i.e., *i* = *j*.

Let bond *i* link atoms *a*_*i*_ and *b*_*i*_ and let bond *j* link atoms *a*_*j*_ and *b*_*j*_. The (*i, j*)th entry of the matrix ***A***(***x, y***) is given by the weighted inner-product

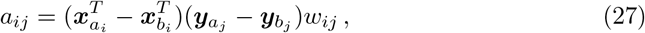

where the weight *w*_*ij*_ is given by

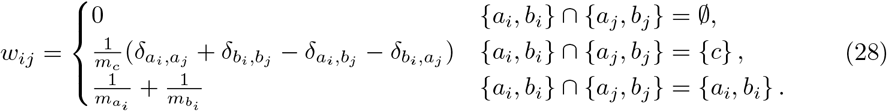

When the matrix ***A***(***x, y***) represents bond constraints for a real molecule, then it is necessarily quite sparse. Consider the row corresponding to the bond between a pair of atoms with valence *r*_1_ and *r*_2_. For this row of the matrix ***A***(***x, y***), there can be at most *r*_1_ + *r*_2_ − 1 nonzero entries, regardless of the overall size of the molecule.

The bond lengths *σ*_*i*_ are normally constant [35] during MD simulations and their values are set by the force fields used.

### 2.5 The importance of solving constraints accurately

Presently, MD simulations usually accept a large relative error when solving the constraint equations. The default value of GROMACS [36] for the maximum allowed relative error for satisfying any constraint with the SHAKE algorithm (`shake tol`) is *τ* = 10^−4^. Other MD packages, like Amber [37], are a bit more demanding (*τ* = 10^−5^) [38]. Notwithstanding, tighter satisfaction of the constraints is sometimes imposed, frequently in simulations performed in the NVE (microcanonical) ensemble [28, 38–48], though also in simulations with a thermostat [28, 49–54] (NVT, NPT). For example, in references [28, 38–54] the authors set a tolerance *τ* (`shake tol`) for the constraints between 10^−7^ and 10^−10^ (except Ref. [41], which uses *τ* = 10^−12^).

Imposing constraints introduces a source of drift in the energy of the analysed system [40, 41], with the size of the drift increasing with the errors in the satisfaction of constraints. One of the principal reasons for performing accurate constraint calculations is to reduce this drift. This is especially important in simulations in the NVE ensemble, where the conservation of the energy is an explicit requisite. Accurate constraints may be also imposed to accurately calculate sought quantities [55, 56] or to avoid undesired effects –like spurious phase transitions [39]–; authors have also reported that improving simulation accuracy eliminated the flying ice cube effect [52].

When conducting simulations using a thermostat, say, in NVT or NPT ensembles, it is not customary to solve the constraint equations accurately, but this is not necessarily the correct approach. Certainly, the energy is not conserved for canonical or NPT ensembles, but there exists a *conserved quantity* (sometimes called *conserved energy*) which is analogous to the conserved energy of the microcanonical (NVE) ensemble. The derivation of the equations of the thermostat (e.g. Nosé-Hoover [57, 58], V-rescale [59]) implies that the conserved quantity is indeed conserved; otherwise, the thermostat equations that are assumed to hold do indeed not hold. There is no reason to believe that the conserved quantity will be conserved if the constraint equations are not solved to the limit of machine precision.

A small drift of the energy does not necessarily indicate that the MD simulation is accurate and reliable. Since different sources of inaccuracy can generate drifts with different signs, large errors in the dynamics can be masked by small drifts resulting from contributions that almost cancel each other. Generally speaking, a large drift of the energy is likely indicating that the dynamics is distorted and the simulation is not reliable, but the converse is not true (a small drift does not necessarily imply that the dynamics is not distorted) [60, 61]. An accurate MD should try to reduce all the sources of errors in the dynamics, hence solve constraints to the maximum affordable accuracy. Otherwise, the calculated quantities will not be reliable. Moreover, small drifts may be misleading, hinting that the simulation is accurate, when it is not.

It is important to appreciate the consequences of solving the constraint equations inaccurately. Let 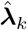 denote the *computed* value of the Lagrange multiplier ***λ***_*k*_; then the *computed* value of ***v***_*k*+1*/*2_ cannot be more accurate than

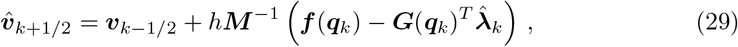

in which case the exact value ***v***_*k*+1*/*2_ satisfies

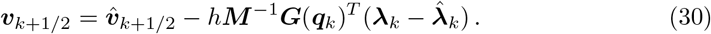

From the equations above we conclude that errors in the calculation of the Lagrange multipliers ***λ***_*n*_ are mathematical equivalent to the actions of an external force, i.e, the term 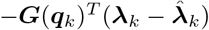. Due to its random nature, this force does not need to be a conservative force. Hence, if the objective is to simulate an *isolated* system, it is critical that we solve the constraint equations accurately.

The distortions of the bond lengths that arise from solving of the constraint equations inaccurately have non-zero average, i.e., the noise is not white, and are highly correlated with their previous values (see results and discussion below and in the S1 File). This is equivalent to using bond lengths which differ from the ones specified by the force fields (whose calibration is highly optimised to accurately describe chemical phenomena), and to using different bond lengths at different times during the simulation. Due to the accumulation of errors throughout many time steps and to the chaotic nature of simulations, the introduced spurious force may distort the dynamics in unpredicted manners, thus reducing the reliability of the calculated quantities. In contrast, if constraints are satisfied allowing a maximum relative error of e.g. 10^−10^ instead of the customary 10^−4^, then the scale of the distortions drops by a factor of 10^6^, see equation (30) and the surrounding paragraph. This significantly reduces the expected impact of spurious forces.

The points stated above indicate that an accurate satisfaction of the constraints is necessary for an appropriate simulation, where the ensemble is respected and the disruptions of the system’s dynamics are reduced.

## 3 ILVES-PC: ILVES for peptide chains

### 3.1 Fundamentals

The only difference between ILVES and SHAKE is how we solve the nonlinear constraint equations; ILVES solves the same system of differential algebraic equations as SHAKE. ILVES uses Newton’s method to solve the nonlinear equations; the involved linear (*linearised*) systems (19) are solved using a direct solver (Gauss-Jordan elimination), which has linear (*O*(*n*), being *n* the number of constraints) scaling due to the sparsity of matrix ***A***. Such sparsity arises from the fact that one atom cannot be covalently bonded to many others. This guarantees the sparsity of ***A*** for biological molecules, and hence makes the ILVES algorithm suitable for constraining all kinds of internal degrees of freedom, including bond and dihedral angles. The ***A*** matrix of relevant bio-polymers can be defined so that it is banded (or nearly banded, with a few nonzero entries outside the band), which enables a special efficiency when solving the system [62]. ILVES-PC relies on a code (*compiled code* [30]) which is specifically designed to be efficient for known structures, such as the amino acid residues that make up peptides and proteins.

### 3.2 Implementation

The implementation of the ILVES algorithm presented in this document, called ILVES-PC, is specifically designed to compute the constraint forces for proteins. Developing code for given types of molecules is the extension to software of a concept which has already proven to be very successful with hardware. The Anton supercomputers were conceived to perform MD simulations of proteins and other biological molecules [9]. Their specific design greatly enhances their performance for these systems compared with general-purpose computers. The algorithm we present in this article also utilises specific features of widely simulated systems in order to increase the performance compared with general-purpose algorithms.

Peptide chains (peptides and proteins) are polymers of variable length formed by repeating blocks of atoms. They are a subset of biological polymers (also including e.g. nucleic acids and polysaccharides), which are just a subset of chemical polymers (which could benefit from the approach presented in this article). Each of the building blocks of a peptide chain (*residues*) has a given connectivity pattern, which is found essentially unchanged in biological molecules (occasionally further atoms can be attached to the protein, e.g. in glycoproteins, or the protein structure can get modified, e.g. at the chromophore of the Green Fluorescent Protein; in addition, hydrogen atoms in carbon rings of side chains can lie in alternative positions). However, 20 given –*proteinogenic*– amino acid connectivities largely dominate the structure of peptides and proteins.

We can make an abstraction of a peptide chain as a graph *G* = (*V, E*). The vertices *V* = {1, 2, 3 …, *n*} represent the atoms and the edges *E ⊆ V* × *V* represent the bonds. Specifically, we have (*a, b*) *∈ E* if and only atoms *a* and *b* are bonded. The graph is undirected because (*a, b*) *∈ E* if and only (*b, a*) *∈ E*. From *G* we can build a new graph *L*(*G*) where each vertex represents a bond and two bonds are connected if and only if they share an atom. The graph *L*(*G*) is known as the line-graph of *G*.

The coefficient matrix of the linear system solved by ILVES-PC, i.e., the matrix ***A*** of equation (19), has the same structure as the adjacency matrix of the bond-graph of the peptide chain. Since the peptide chain is composed of less than 30 different building blocks, the main features of the matrix ***A*** can be described using less than 30 different submatrices. These matrices correspond to the proteinogenic amino acids, some of them having slightly different configurations due to different locations of hydrogen atoms. In truth, we need a few more subroutines to account for, say, the beginning and the end of the chain. We have written subroutines for solving the linear subsystems corresponding to each of these submatrices. To solve the entire linear system, we iterate over the subsystems of the linear system, calling the required subroutine. We ensure that submatrices corresponding to identical amino acids have the same structure by always numbering the bonds of each amino acid in the same order. This allows us to generate loop-free subroutines that store the relevant data contiguously in memory and do direct memory access instead of relying on the auxiliary data structures and the indirect memory access patterns that are so typical of direct solvers for sparse linear systems. These ideas are all further adaptations of the compiled code approach utilised in [30]. By choosing the bond numbering of each amino acid we can reduce the fill-in during the factorization of the matrix. Fill-in are entries that are exactly zero in the original matrix, but become non-zero during the factorization process. We can work with peptide chains with any bond numbering. Before simulating a new protein for the very first time, we first explore the topology to identify the individual amino acid residues. Then, we renumber the bonds to match the numbering required by our specific implementation. The structural information can be saved and recycled if further simulations are required.

ILVES-PC exploits modern processors’ computational resources by assigning a subset of amino acids of a peptide chain to different hardware threads. We ensure that each thread is assigned a similar number of matrix elements. To concurrently apply Gauss-Jordan elimination to the submatrices corresponding to different amino acids, we rely on the Schur complement method [63].

Consider the example presented in Fig 1. We want to make the elimination at the banded coordinate matrix of (a) –which exemplifies ***A***– using three threads (magenta, purple, and yellow). First (b), each of the threads works with its own *private* submatrix, trying to fill the lower left-hand (subdiagonal) corner with zeros and the diagonal with ones. The threads also update the shared (blue) rows using mutual exclusion (mutex) locks. They repeat this step (c) in the upper-hand corner of their submatrix. These two steps may produce fill-ins. Then, the master thread processes the submatrix comprised of the shared (blue) rows (d), while the other two threads wait until this step is completed. Finally (e), all threads work in parallel to clean the fill-ins produced in steps (b) and (c). In the case of ILVES-PC, the thread private data corresponds to constraints within a given amino acid, while the shared rows correspond to the constraint that connects two amino acids (which corresponds to a peptide bond).

**Fig 1.**
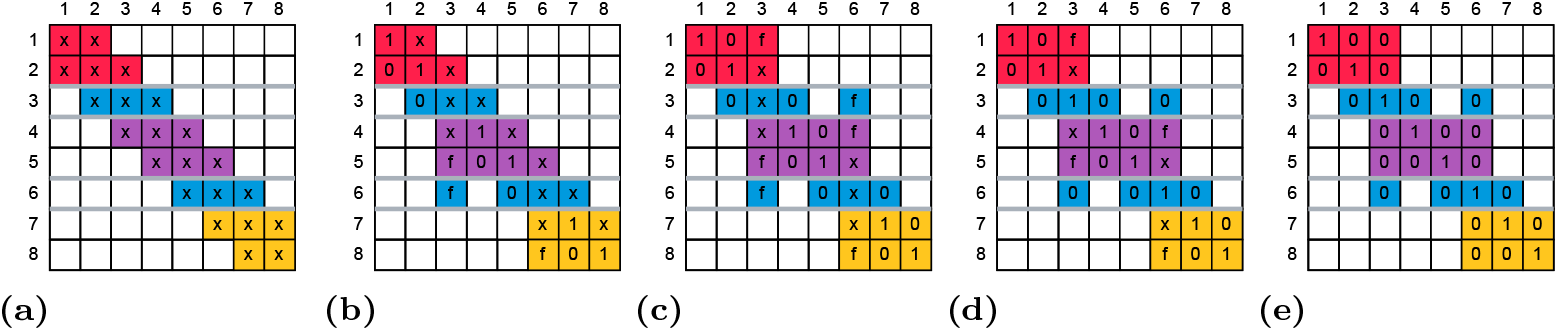
Steps to apply Gauss-Jordan elimination to a banded matrix in parallel using three threads via the Schur complement method. The entries of the matrix are represented with and *x* and the fill-ins with an *f*. The magenta, purple and yellow entries are private to threads, while blue entries are shared between threads.

While a general-purpose implementation of ILVES would almost certainly rely mainly on coarse-grain synchronization mechanisms such as thread barriers, the precise knowledge of the structure of the linear system corresponding to proteins allows us to use very fine-grain synchronization mechanisms. We use lightweight mutex locks to protect the shared rows of the linear system from data races, so that each mutex involves only a pair of threads. Similarly, during the update phase at the end of each Newton step, the positions and velocities of each atom are protected by individual mutex locks.

## 4 Materials and methods

We have integrated ILVES-PC into the popular GROMACS molecular simulation package [36]. Our solver can be used as an alternative to SHAKE and P-LINCS when solving the bond constraints of proteins (only consisting of amino acid residues) without disulfide bonds [64]. In addition we have extended the code of P-LINCS to accept a tolerance *τ >* 0 for the satisfaction constraints on all bonds (all-bonds), as in SHAKE and ILVES-PC, so that the three solvers can be compared on an equal footing. All the tested algorithms iterate until

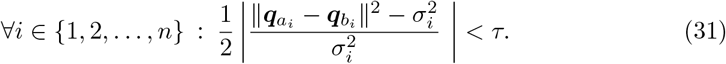

where the *i*th bond is between atoms *a*_*i*_ and *b*_*i*_. It is straightforward to verify that

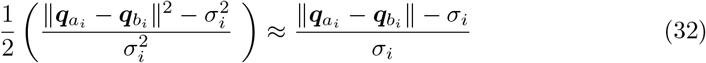

is a good approximation when the *i*th constraint equation is almost satisfied, i.e., when 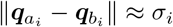 is a good approximation. It follows that *τ* is a good approximation of an upper bound for the largest relative error for the bond lengths.

As already mentioned, P-LINCS uses a truncated Neumann series to approximate the solution; then P-LINCS applies an iterative correction phase. The accuracy of the expansion is determined by its order (`lincs order`), and the accuracy of the correction phase is determined by the number of iterations (`lincs iter`). Both the order of the expansion and the number of iterations of the correction phase are set at GROMACS’ startup and do not change throughout a given simulation. With our modification, P-LINCS keeps iterating in the correcting phase until all constraints are solved with the specified tolerance (`shake tol`). We found that the additional calculations due to this modification (i.e. the calculations to check the degree of constraint satisfaction) typically increase P-LINCS’ execution time by 4% to 9%.

### 4.1 Experimental setup

For the experiments carried out in this work we have used our modified version of GROMACS 2020.1 in double precision (`-DGMX DOUBLE=on`) compiled using GCC-10.1.0. All simulations were performed on a computer with two Intel Xeon Platinum 8160 CPUs. Each processors has 24 physical cores. The main characteristics of our computer are presented in Table 1. We have used a GROMACS OpenMP version for multi-thread executions.

**Table 1.** Main features of our computer.

|  |  |
| --- | --- |
| Processor | 2 × Intel Xeon Platinum 8160 |
| Cores | 2 × 24 |
| AVX-512 FMA units | 2 (per core) |
| L1 cache (I, D) | 8-way 32 KiB (per core) |
| L2 cache | 16-way 1024 KiB (per core) |
| LLC | 11-way 33 MiB (shared) |
| Main Memory | 96 GiB DDR4 (12 × 8 GB 2667 MHz DIMM), 6 channels |
| OS | SUSE Linux Enterprise Server 12 SP2, kernel 4.4.120-92.70-default |

### 4.2 Simulations

Four proteins have been simulated in this study, namely, ubiquitin, COVID-19 main protease, human SSU processome and barnase. A summary of their features can be found in Table 2. In order to assess the performance of the protein-specific constraint solver (ILVES-PC) presented in this article, simulations of the first three proteins were performed. They cover a wide range of protein sizes: 76 to 2722 amino acid residues; SSU processome is unusually large, and was mainly chosen for testing parallel scalability.

**Table 2.** Proteins simulated in this article. The number of amino acid residues, atoms and constraints (all-bonds), the PDB codes and articles of reference are included.

| Name | # resid. | # atoms | # constr. | Code | Reference |
| --- | --- | --- | --- | --- | --- |
| Ubiquitin | 76 | 1231 | 1237 | 1UBQ | [65] |
| COVID-19<br>main protease | 304 | 4645 | 4981 | 5R7Y | [66] |
| Human SSU<br>processome | 2722 | 40802 | 41209 | 7MQA<br>bundle 3<br>chain F | [67] |
| Barnase | 110 | 1700 | 1721 | 1A2P | [68] |

The atomic coordinates of these proteins were taken from the RCSB Protein Data Bank (www.rcsb.org, see the PDB codes in Table 2).

Our modified version of GROMACS was used to run and analyse the simulations, which were set up with the CHARMM36 force field including CGenFF version 4.1 (last update on March 28, 2019) [69]. The ionizable residues of selected proteins were protonated in all the cases as default (pH 7) in GROMACS, and after adding explicitly TIP3P water molecules [70], chloride or sodium counterions were added to neutralize the systems. A truncated dodecahedral was chosen as the simulation box, and the minimum distance between the protein and the box edge was set to 1 nm. Periodic boundary conditions (PBC) were imposed. A minimization phase of the solvated systems was performed (maximum of 20000 steps) with the steepest descent algorithm [71]. One (for the evaluation of performance with ubiquitin, COVID-19 main protease and human SSU processome) or three (for the evaluation of accuracy with barnase and ubiquitin) simulation replicas were launched under each setting, enabling the averaging of results in the latter case. Systems were gradually heated through a heating ladder that allowed to increase the temperature tier by tier in a number of NVT steps (50 K every 50 ns; 1 fs time step) until reaching the target temperature (either 298 or 400 K [72]). Three consecutive steps for system equilibration followed. The first one (NVT) ran for 100 ps with restraints imposed on the heavy atoms (protein and water) and the V-rescale thermostat [59] used to keep the targeted temperature (coupling strength parameter *τ*_*T*_ = 0.1 ps). The second equilibration step (NPT) ran for 100 ps (2 fs time step) without any restraint, with the V-rescale thermostat (*τ*_*T*_ = 0.1 ps) [59] and the Berendsen barostat used to set the pressure to 1 atm (coupling strength parameter *τ*_*p*_ = 2.0 ps) [73]. The third equilibration step (NPT) ran for 200 ps (2 fs time step) using the V-rescale thermostat (*τ*_*T*_ = 0.1 ps) and the Parrinello-Rahman barostat [74] (1 atm; *τ*_*p*_ = 2.0 ps). The configuration reached after these steps was the starting point for different production runs carried out in either the NPT or NVE thermodynamic ensembles.

For calculations of performance (NPT) and accuracy (NPT or NVE), the thermostat, barostat and the rest of parameters were set up in the production phase as done in the third equilibration step, except for the NVE simulations (used only for ubiquitin and barnase), where neither thermostat nor barostat were used. Production runs launched for calculation of performance consisted in 50k steps (2 fs time step for ubiquitin and COVID-19 main protease) or 10k steps (2 fs time step for human SSU-processome). On the other hand, production runs for calculation of the simulations accuracy (energy drift analysis of barnase and ubiquitin) consisted of half a million (5 *·* 10^5^) steps (2 fs time step, for a total simulation time of 10 ns). A record of the conserved energy values (in NPT simulations) or of the total energy (in NVE simulations) at every 1000 steps (0.2 ps) was obtained. These numbers were used to calculate the energy drift.

As general parameters of the simulations the Verlet cutoff-scheme algorithm [75, 76] was used for van der Waals interactions and the PME method [77] for electrostatic interactions, both with a radius cutoff of 1.2 nm as recommended by the authors of the CHARMM36 force field [69]. A radius cutoff of 1.0 nm was also set in simulations used for performance calculations. Velocities correction due to the thermostat were done every 10 timesteps (default values of GROMACS).

For the tests performed to assess simulation performance, tolerance values (*τ*) of 10^−4^, 10^−6^, 10^−8^, 10^−10^ and 10^−12^ have been used to constraint all bonds. The constraint algorithms tested include SHAKE, the modified version of P-LINCS and ILVES-PC. In the case of P-LINCS also the `lincs_order` parameter has been tested and values of 4 and 8 are combined with the tolerance values listed above. In simulations carried out to analyse the accuracy the SHAKE algorithm was tested with constraints both on all-bonds and on bonds connecting with hydrogen atoms (H-bonds; results obtained with these constraints appear in Fig 1 of the S1 File). Tolerance (*τ*) values of 3.1423 *·* 10^−4^, 10^−4^, 3.1423 *·* 10^−5^, 10^−5^, 10^−6^, 10^−7^, 10^−8^, and 10^−10^ have been tested.

The analysis of the energy drift has been done by taking also into account the Verlet buffer size for the pair-list neighboring search. The pair-list neighboring search is considered one of the two most important sources of energy drift in MD simulations (the other one being that related to the constraints algorithm). The GROMACS parameter used to establish the Verlet buffer size (`Verlet-buffer-tolerance`) was set here to 5 *·* 10^−5^ kJ/(mol *·* ps) to permit a lower level of drift than that allowed for GROMACS’ default value (5 *·* 10^−3^ kJ/(mol *·* ps)). This way, it is possible to display the drift effect due to constraints more clearly. Data presented at the Fig 1 of the S1 File also include results obtained for a `Verlet-buffer-tolerance` of 5 *·* 10^−3^ kJ/(mol *·* ps).

## 5 Results and discussion

In this section, results of the test calculations are summarised. In the first subsection, an analysis of the reliability of the simulation as a function of the accuracy in satisfying constraints is presented, whereas an analysis of the efficiency (performance) of the calculations is displayed in the second subsection.

### 5.1 Physics

One of our goals is to understand the connection between the energy drift in MD simulations of proteins and the tolerance *τ* of the constraint algorithm (see equation (31) for the definition of *τ*). For this purpose, we have simulated ubiquitin and barnase in the NVE and NPT ensembles (with a V-rescale thermostat in the latter), using SHAKE to impose constraints. We choose SHAKE because the slow and steady convergence of this algorithm ensures that the largest relative error for the bond lengths is essentially equal to the tolerance *τ* of the constraint algorithm. The time evolution of the energy followed regular straight lines in large enough time scales; hence we calculated the drift as the slope of the energy-vs-time relationship [60] divided by the number of degrees of freedom of the system *N*_*df*_. We defined the latter as: *N*_*idf*_ = 3*m* − *n* − 6, where *m* is the number of atoms in the protein, *n* is the number of imposed constraints, and the −6 term accounts for rotations and translations. In the NPT simulations the temperature was set to 298 and to 400 K (this latter only for barnase), whereas the pressure was set to 1 atmosphere. To convert Joule per mol to units of *k*_*B*_*T* we have divided by 2477.7 if T=298 K (it includes the NVE simulations, where no T is imposed during the production stage, but 298 K was initially set), and divided by 3325.8 if T=400 K. The time step used for all the simulations was 2 fs.

The results of the drift calculations are presented in Fig 2 [79]. It is observed that the energy drift decays rapidly as the tolerance *τ* is reduced. The results presented in Fig 2 are consistent with the literature, where *τ* = 10^−7^ is used to reduce the energy drift [47–49] to acceptable levels. Fig 2 suggests that one should choose a constraint tolerance *τ* which is at least as small at *τ* = 10^−6^.

**Fig 2.**
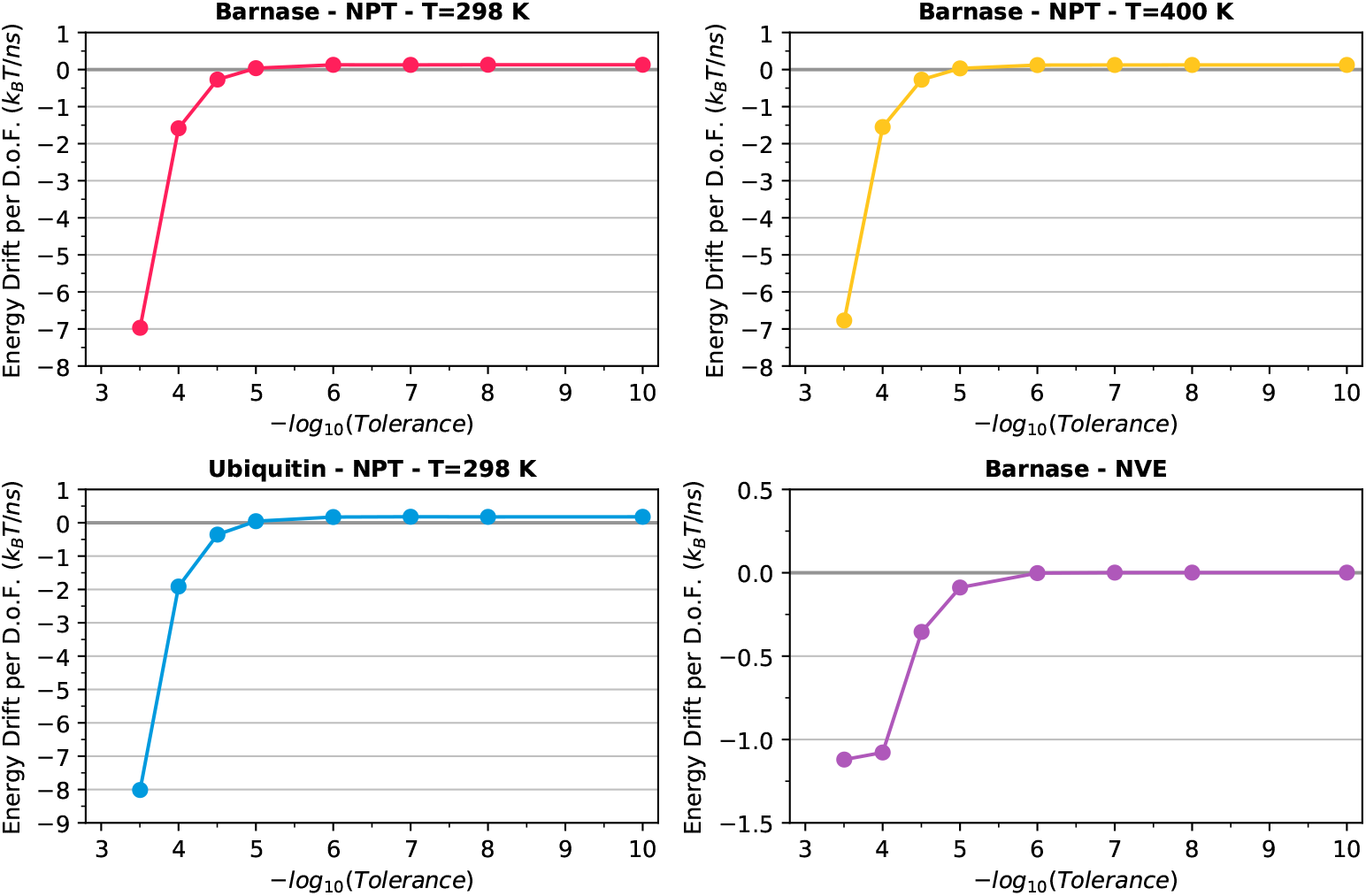
Drifts of the conserved energy of the thermostat (NPT ensemble) and of the energy (NVE ensemble) per degree of freedom as a function of the tolerance in satisfying the constraints. Top: barnase in the NPT ensemble at T=298 K (left) and T=400 K (right); Bottom: ubiquitin in the NPT ensemble at T=298 K (left) and barnase in the NVE ensemble (right).

As a further evidence of the importance of using small values of *τ*, in Fig 3 it is observed that higher tolerances in NVE simulations led to significant drift of the initial temperature set for the simulations. Conversely, if we use *τ ≤* 10^−6^ the temperature is approximately preserved over the analysed time range. Other authors have also found that the low accuracy in solving the constraints gives rise to inaccurate temperatures, and that the default parameters of LINCS lead to non-converged results [78].

**Fig 3.**
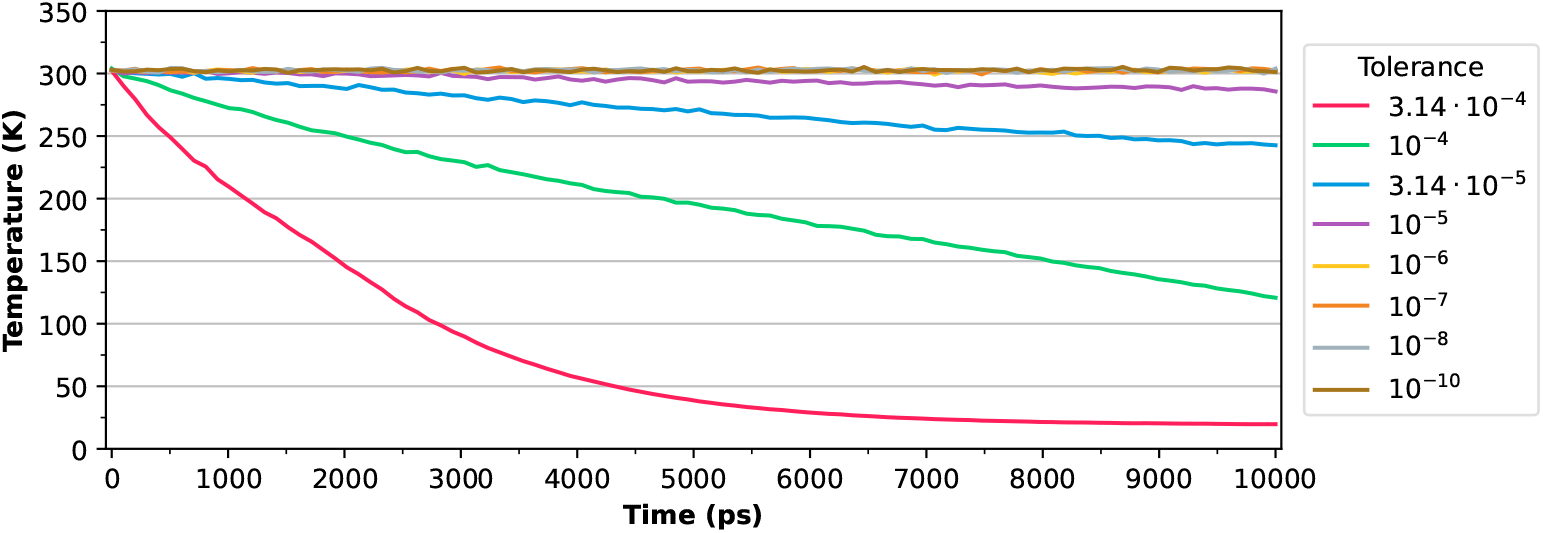
Temperature evolution over time of barnase in the NVE ensemble using different tolerances in satisfying the constraints.

The non-negligible drifts presented in Fig 2 suggest that the mechanics of the whole molecule have distorted. Here, the dynamics has been examined on a finer level by measuring the actual bond lengths, whose value is expected to be frozen since constraints on all bonds are imposed, as a function of time. Our primary goal is to answer the questions: *Do the distortions of the bond lengths have zero average? Are the distortions of the bond lengths time-correlated?*

To delve into these questions, an analysis of bond length error (actual *vs*. expected value) has been performed for all the constrained bonds in our simulations. Plots at the top and center panels of Fig 4 are displayed as an example and correspond to 2000 time steps (1 time step = 2 fs) from an NPT production trajectory of ubiquitin run with the SHAKE constraint solver with tolerances of *τ* = 10^−4^ and *τ* = 10^−10^, respectively.

**Fig 4.**
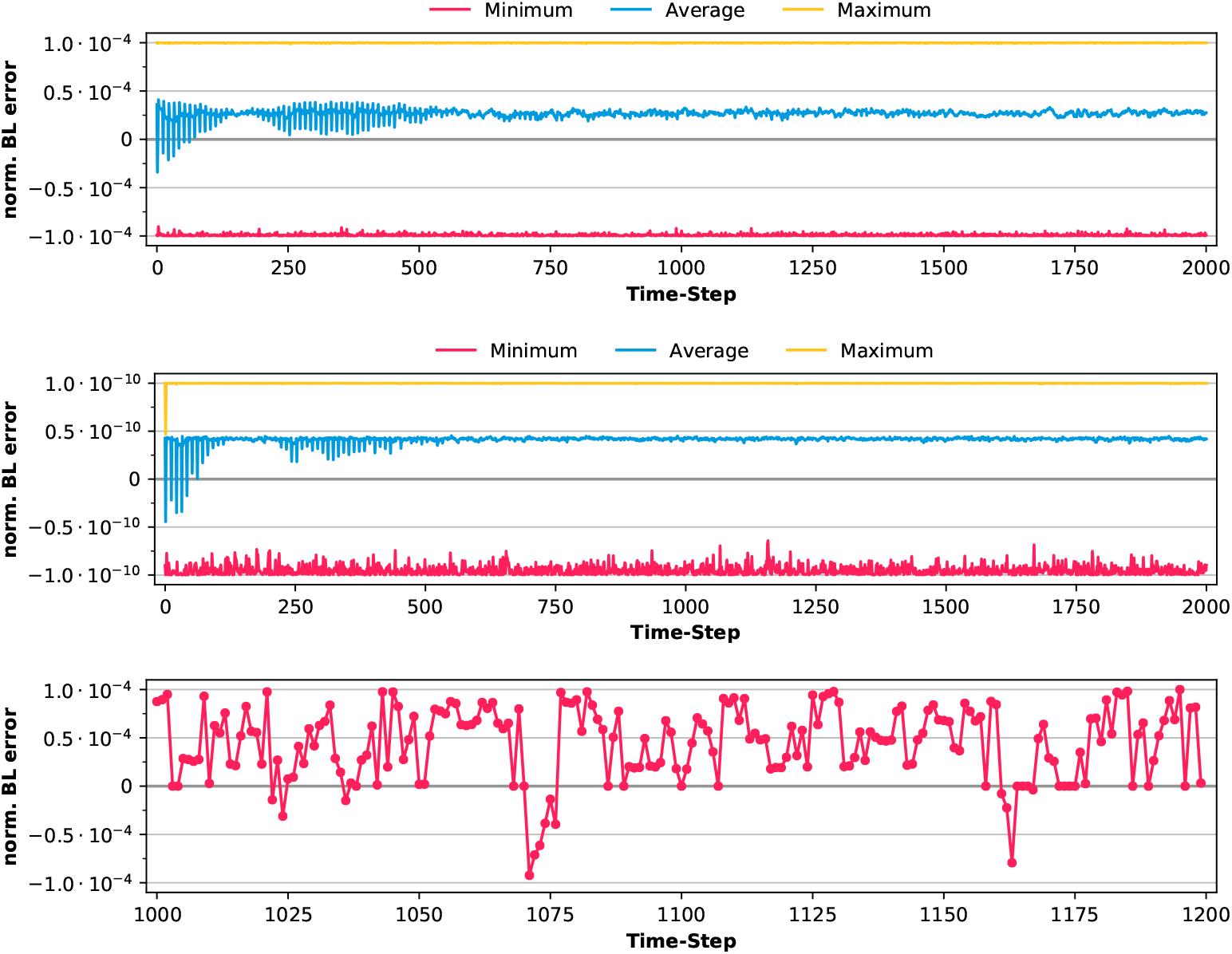
Normalised error over time for the bond lengths in MD runs of ubiquitin with SHAKE. The top and central panels show the average, maximum and minimum values of the normalised error for all the constrained bonds (all-bonds). The top panel corresponds to the tolerance of *τ* = 10^−4^ while the central one corresponds to *τ* = 10^−10^. The bottom panel shows a representative normalised error profile obtained for a constrained bond.

These plots include the *average* 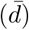, *maximum* (*d*_Max_) and *minimum* (*d*_min_) bond length errors over time obtained for all the constrained bonds. These parameters are defined as:

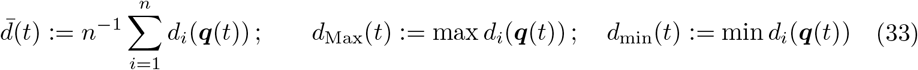

where *d*_*i*_ is the relative error for the *i*th bond length, i.e.,

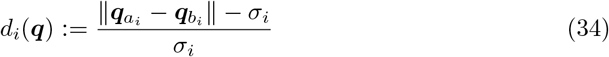

and *n* is the number of constraints.

The maximum and minimum values, *d*_Max_ and *d*_min_ (yellow and red lines, respectively), are approximately the established value for the tolerance of SHAKE, which indicates that the solver usually does not provide solutions much more accurate than that required for the user [80]. The average 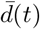 (blue line) is positive along the time, which implies that the bond lengths are, on average, longer than expected. Qualitatively, the patterns of the error plots persist if the tolerance of the constraint solver is changed, e.g. to *τ* = 10^−10^ (central panel in Fig 4). However, as expected, the size of the distortions is six orders of magnitude lower than those obtained for *τ* = 10^−4^ (top panel in Fig 4). This element confirms that a tighter tolerance for satisfying the constraints largely reduces the bond length distortions in the simulated system.

The bond length distortions presented in the plots at the top and central panels of Fig 4 show a Pearson correlation with their immediately previous value of 0.41 *±* 0.12 (average *±* SD, see Fig 15 of the S1 File).

Similar plots have been obtained for one of the simulations of ubiquitin run with the P-LINCS solver (`lincs_order`=4, `lincs_iter`=2, and the rest of parameters identical to those used in the simulation presented in Fig 4) and presented in Fig 5 of the SI File. The normalised bond length error plot presenting the largest values among all the constrained bonds is depicted in the top panel of SI Fig 5. The average distortion (bottom panel) obtained over time for all the constrained bonds (blue line) with P-LINCS mounts to 2.17 *·* 10^−5^, a lower value than that obtained for ubiquitin simulated with SHAKE (2.67 *·* 10^−5^). An interesting element one can point out from this analysis is that the bond length error profiles obtained with SHAKE present much sharper and chaotic peaks than those obtained with P-LINCS. As a result of this, a higher temporal correlation between the normalised errors and their previous value is observed with P-LINCS.

**Fig 5.**
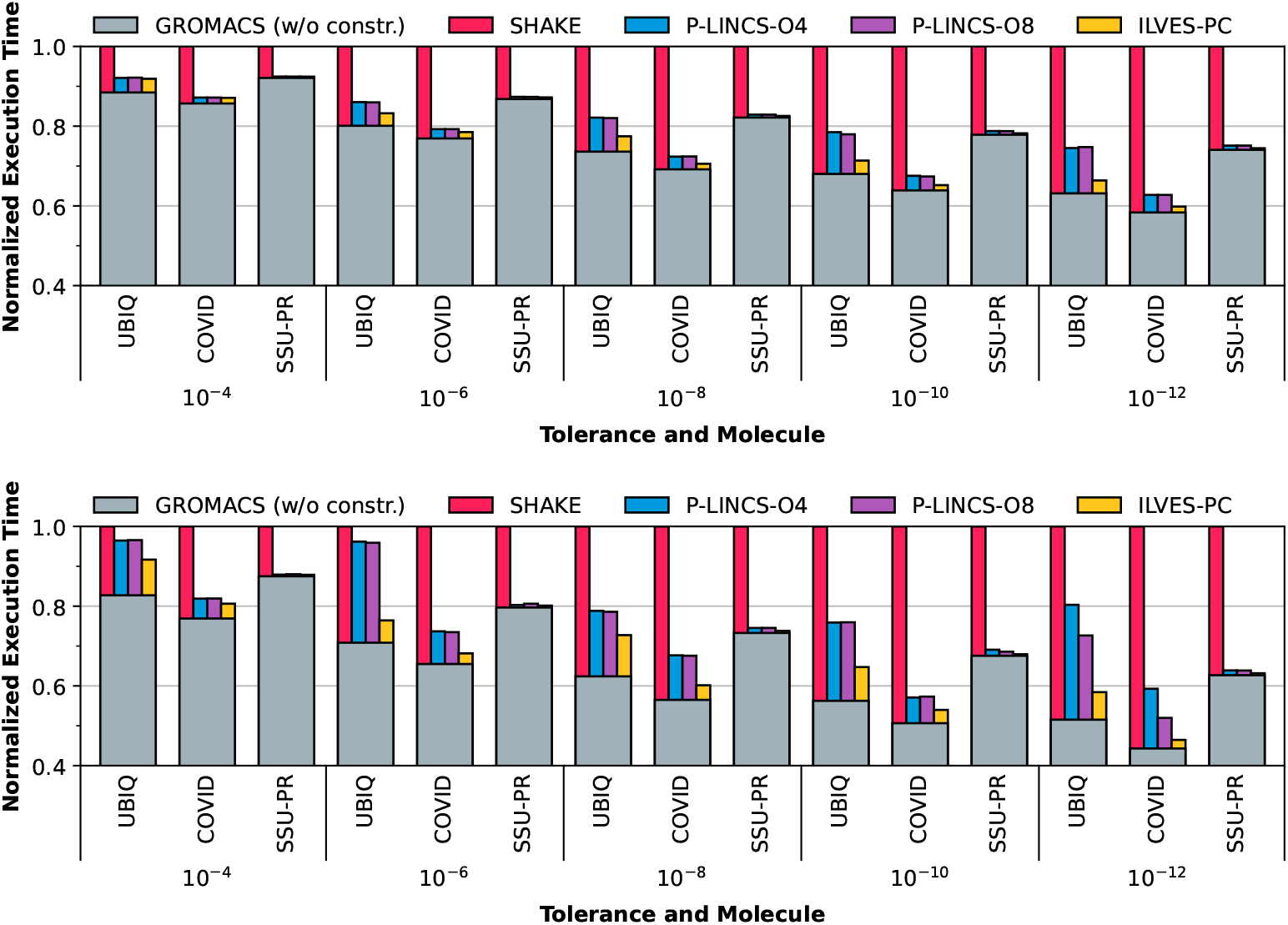
Normalized execution times for different accuracy limits (tolerances) and three different proteins. The height of the thick grey bars represents the execution time of GROMACS excluding the constraints computations. The height of the magenta, blue, purple, and yellow bars represent the time required by the different constraint solvers. Top: parallel execution with 24 threads; bottom: ibid. with 48 threads. Note that the *y*-axis starts at *y* = 0.4 rather than *y* = 0. This has been done to emphasize the differences between the constraint solvers.

The bottom panel of Fig 4 shows a representative normalised bond length error profile for a given constrained bond. The most important feature one can notice there is that the error is almost exclusively positive. This the most recurrent trend in all the bonds, see Figs 7 to 14 of the S1 File. Some other examples of this plot –grouped per bonds of the same nature– are presented in Figs 7 to 14 of the S1 File, and additional details thereof are pointed out there.

### 5.2 Performance

We have evaluated the performance of the state-of-the-art constraint solvers (SHAKE and P-LINCS) and of the Newton method-based solver presented in this article (ILVES-PC), imposing constraints on all bond lengths. The modified version of P-LINCS (which stops iterating when the largest relative error for the bond lengths is less than the specified tolerance *τ*) was used, and two values of its matrix expansion parameter (`lincs order`) were assessed: 4 (the default) and 8 (a commonly used value). Results obtained for these tests are labeled in the figures as P-LINCS-O4 and P-LINCS-O8, respectively. Other input parameters of the simulations were kept at their default values in GROMACS. Simulations were carried out for ubiquitin, COVID-19 main protease and human SSU processome in the NPT ensemble (298 K and 1 atm). The results are presented in Figs 5, 6, 7, and 8.

**Fig 6.**
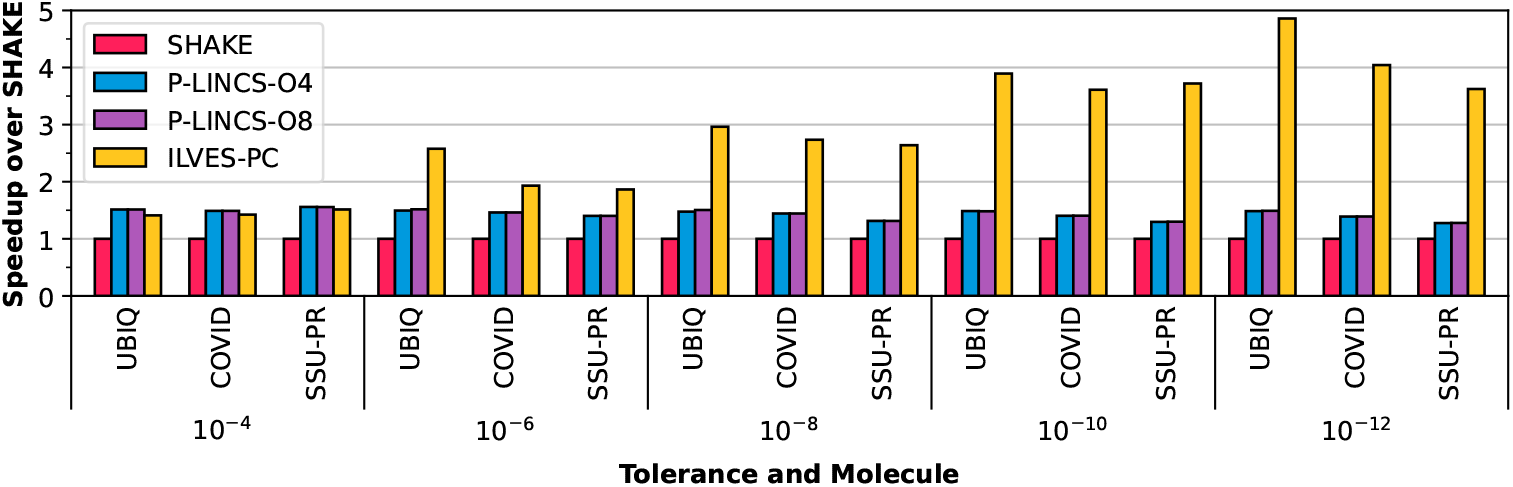
Single-thread speedup over the SHAKE algorithm of ILVES-PC and P-LINCS for different tolerances and test cases.

**Fig 7.**
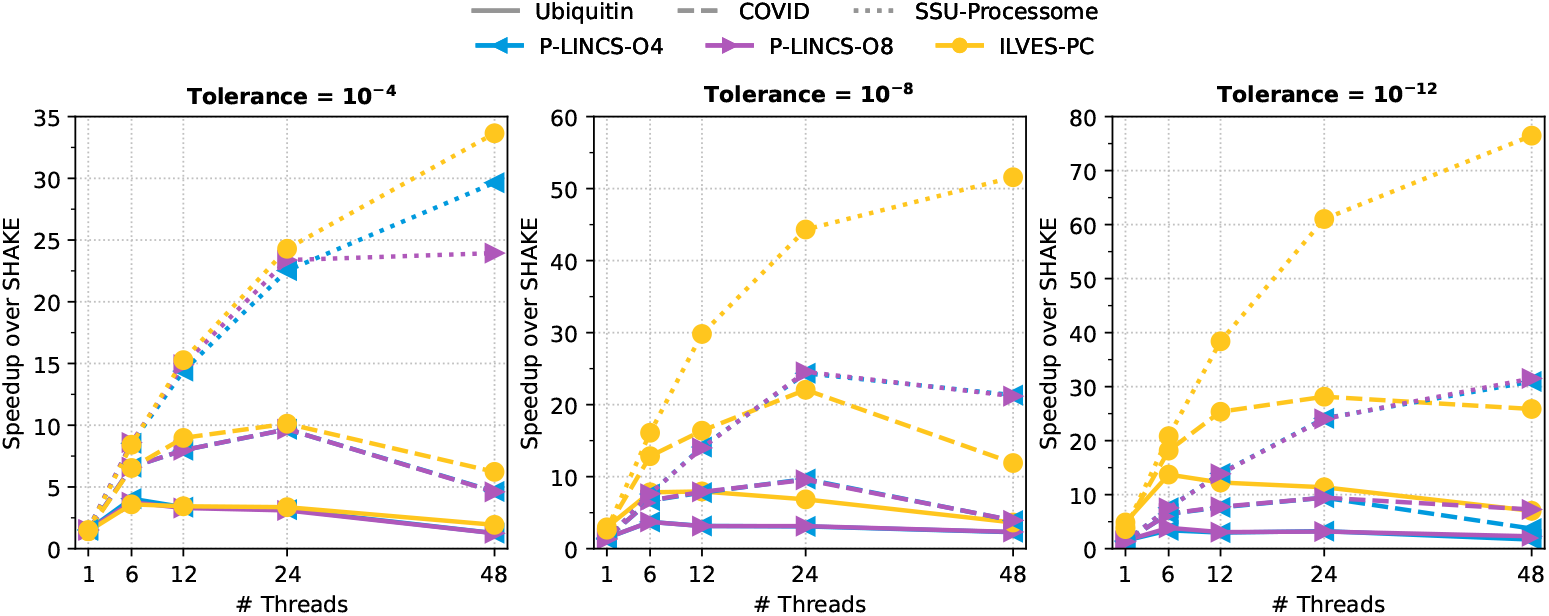
Multi-threaded speedup of ILVES-PC and P-LINCS over the SHAKE algorithm for different tolerances and test cases. Note the different y-axis scale of each plot.

**Fig 8.**
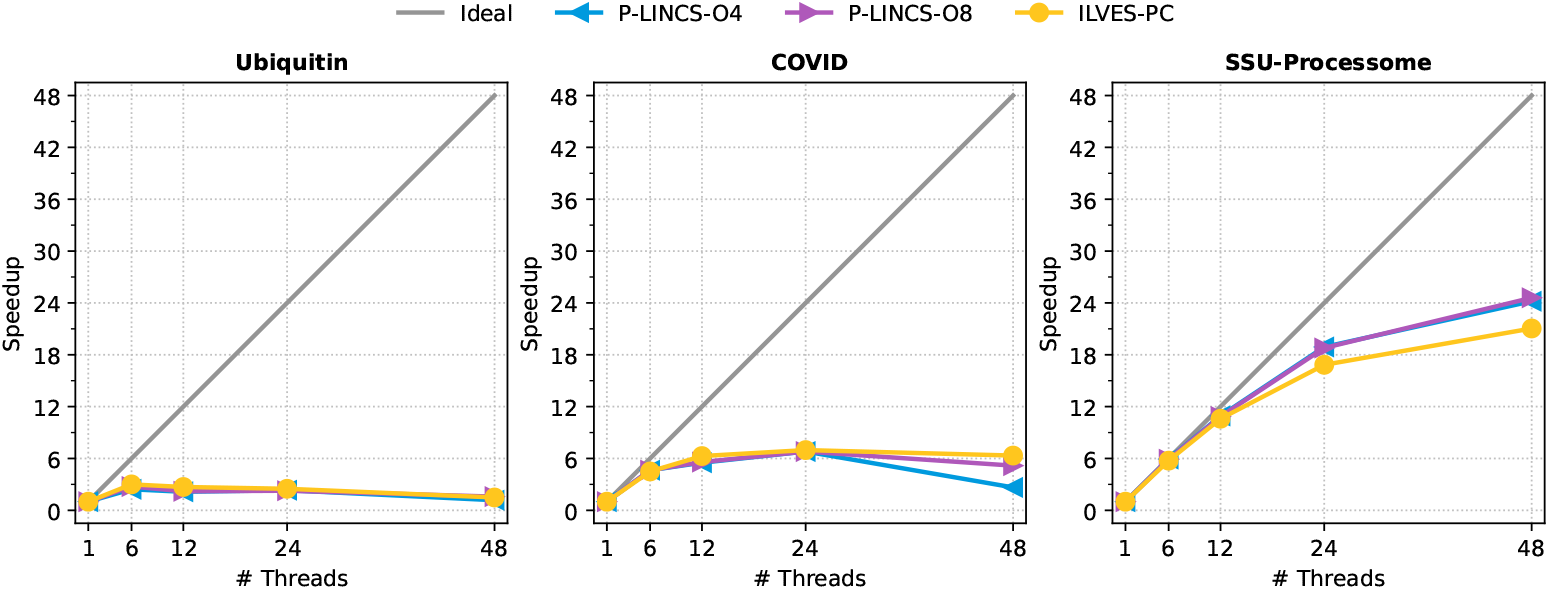
Speedup over serial execution of P-LINCS and ILVES-PC for different test cases and numbers of threads.

Fig 5 presents the total (parallel, wall-clock) execution time. Stacked bars show the fraction of the time which is spent solving the constraints imposed on the protein (not on the solvent) in the production phase of the simulations and the rest of the execution time (gray background) [79]. The text labels (UBIQ, COVID and SSU-PR) along the *x*-axis of Fig. 5 represent the three analysed proteins (ubiquitin, COVID-19 main protease and the –atypically large– human SSU processome, respectively). The numerical labels along the *x*-axis represent the largest acceptable relative error for a constraint in each simulations, i.e., *τ* = 10^−4^, 10^−6^,…, 10^−12^.

The value *τ* = 10^−4^ is the GROMACS default value; we omit larger values of *τ* because, as presented in the previous section, they produce result that do not converge [81]. We found that *τ* = 10^−12^ is near the lowest relative error that can be consistently achieved during our simulations. This should not be perceived as general result as this value depends on the specific equations being solved and the floating point number system used.

The results presented in Fig 5 indicate that the performance of SHAKE, whose implementations are most commonly serial –in contrast to P-LINCS and ILVES-PC–, is the worst among the analysed solvers. This implies that SHAKE requires higher ratios of the total execution time, which is aggravated by increasing the number of threads. This is a natural consequence of Amdahl’s Law [7], that is, the performance improvement obtained by parallel execution is limited by the sequential fraction of the application. This problem is likely to be exacerbated by the continuous increase in the number of cores in the processors. ILVES-PC performs better than the state-of-the-art in nearly all the analysed cases, and its relative advantage increases as higher accuracy is demanded. Moreover, since the computation time is similar for high accuracy (e.g. *τ* = 10^−12^) and low accuracy (e.g. *τ* = 10^−4^), accurate calculations become affordable using ILVES-PC.

It is entirely possible to view Fig 5 and conclude that it is a waste of time to improve the quality of parallel constraint solvers. After all, the majority of the execution time is spent outside the constraint solver, so why should we bother improving the constraint solver? This line of reasoning ignores two key points:

1. It is easy to question the conclusions drawn from inaccurate simulations that show a significant violation of the fundamental principle of conservation of energy. High accuracy is costly unless the constraint solver is parallel.
2. The efficient use of large computers require algorithms that scale well. It is wasteful to assigning 48 cores to a problem if the speedup is only 4. This precludes the use of algorithms that scale poorly.

Fig 6 shows the single-threaded speedup of P-LINCS and ILVES-PC compared with SHAKE [79]. Both P-LINCS and ILVES-PC show a speedup of approximately 1.30 over SHAKE using the GROMACS default tolerance (*τ* = 10^−4^). While P-LINCS preserves a speedup over SHAKE of between 1.25 and 1.5 for the whole range of tolerances, ILVES-PC’s speedup with respect to both SHAKE and P-LINCS increases as the tolerance *τ* decreases, achieving an average speedup over SHAKE of 4.2 at the smallest value of *τ*.

We have also executed P-LINCS and ILVES-PC using different thread counts. Fig 7 presents the multi-threaded speedup of P-LINCS and ILVES-PC over SHAKE for three different tolerances. The parallel speedups over SHAKE of both P-LINCS and ILVES are similar for all thread counts using GROMACS’ default tolerance (*τ* = 10^−4^), achieving maximum speedups over SHAKE of 4×, 10×, and 34× for the three executed test cases. As we decrease the tolerance, ILVES-PC speedups over SHAKE dramatically increase while P-LINCS shows similar speedups for the whole range of tolerances. At the minimum tolerance (*τ* = 10^−12^), ILVES-PC achieves maximum speedups over SHAKE of 13×, 28×, and 76× for the three analyzed proteins. This is not surprising. ILVES-PC is based on Newton’s method which normally has quadratic convergence, while there is no reason to expect more than linear convergence from P-LINCS.

Fig 8 shows the parallel scalability of P-LINCS and ILVES using different numbers of threads setting the tolerance to *τ* = 10^−12^ [79]. Nearly identical results were obtained for the whole range of tolerances. The scalability of both solvers is similar for all three test cases. ILVES-PC performs slightly better than P-LINCS for the two representative proteins. P-LINCS and ILVES-PC require regular synchronization between threads, and the size of their parallel tasks depends on the number of bonds of the molecule. Hence the size of the test case is important for scalability. If the molecule is sufficiently small and the number of cores is sufficiently large, then the parallel overhead will dominate and no solver can be efficient. Conversely, if the molecule is sufficiently large compared with the number of cores, then good parallel performance is not theoretically impossible.

## 6 Conclusions and future work

The constraint solver introduced in this article demonstrates that it is possible to conduct efficient parallel simulations of polymers by utilising the chemical structure. We have also presented arguments supporting that constraint equations must be solved accurately, including the fact that the default configuration of popular MD packages leads to situations where the constraint solver does not lead to converged results.

The introduced algorithm is well-suited for accurate calculations. It is faster than state-of-the-art constraint algorithms for most of the analysed cases and equally fast for all other cases. The use of Newton’s method ensures that we can reach high accuracy with a small increase in the computational effort compared with low accuracy simulations.

Future stages of this project include tackling the case of imposing constraints on H-bonds, the use of inexact Newton methods –such as symmetric approximations of the Jacobian– in order to achieve higher computational efficiency, the extension of ILVES to nucleic acids (ILVES-NA), MPI parallelization, SIMD vectorization as well as a general version of ILVES which can calculate constraints in every molecule.

## Acknowledgments

We want to thank Jesús Asín (Universidad de Zaragoza) and Benjamin Dalton (Freie Universität Berlin) for our interesting discussions.

## Supporting Information

**S1File**. Supporting Information of *Accurate and efficient constrained molecular dynamics of polymers through Newton’s method and special purpose code*.

## Notes

### Competing Interest Statement

The authors have declared no competing interest.

### Summary of Updates

Supporting information file added.

