## Supporting Information File. for "Accurate and efficient constrained molecular dynamics of polymers through Newton’s method and special purpose code"

[9]*Independent scholar*

(Dated: September 27, 2022)

#### Abstract

In this document we present data that complements the results displayed in the main article. Such data is related to the energy drift and the evolution of the temperature, errors in the bond lengths, the frequency of the crashes in the simulations and the performance of the software.



### I. RESULTS - PHYSICS

#### A. Energy variations

The data included in Table I corresponds to the graphs of the drift per degree of freedom presented in the main article (Fig 2). In Table II the drifts per degree of freedom are summarised for simulations where only constraints on bonds connecting hydrogen atoms (H-bonds) were imposed. In these cases, the number of constraints imposed on ubiquitin and barnase was 629 and 836, respectively, which has an impact on the degrees of freedom used for the calculation. Fig 1 displays the data presented on Table II. A comparison of the data displayed in Tables I and II shows a lower drift-per-degree of freedom (on average) if constraints are imposed on all bonds. For high accuracy in satisfying the constraints (tolerance $10^{-10}$) the average drift for Ubiq-NPT, Bar-NPT and Bar-NVE is 0.28 $k_BT/ns$ if constraints are imposed only on H-bonds, but is 0.10 $k_BT/ns$ if constraints are imposed on all bonds.

TABLE I: Drifts of the energy (NVE ensemble) and of the conserved energy of the thermostat (NPT ensemble) per degree of freedom obtained in simulations of barnase and ubiquitin with several tolerances applied to constrain all bonds. All drifts in units of $k_BT/ns$.

| Molecule: | barnase | barnase | barnase | ubiquitin |
|---|---|---|---|---|
| Ensemble: | NVE | NPT | NPT | NPT |
| T (K) | - | 298 | 400 | 298 |
| Tolerance | | | | |
| $3.14{\cdot}10^{-4}$ | -1.120 | -6.969 | -6.771 | -8.014 |
| $10^{-4}$ | -1.078 | -1.585 | -1.550 | -1.912 |
| $3.14{\cdot}10^{-5}$ | -0.355 | -0.269 | -0.273 | -0.355 |
| $10^{-5}$ | -0.088 | 0.038 | 0.032 | 0.047 |
| $10^{-6}$ | -0.002 | 0.128 | 0.121 | 0.171 |
| $10^{-7}$ | 0.001 | 0.127 | 0.124 | 0.179 |
| $10^{-8}$ | 0.001 | 0.130 | 0.125 | 0.176 |
| $10^{-10}$ | 0.001 | 0.130 | 0.125 | 0.178 |

TABLE II: Drifts of the energy (NVE ensemble) and of the conserved energy of the thermostat (NPT ensemble) per degree of freedom obtained in simulations of barnase and ubiquitin with several tolerances applied to constrain H-bonds. All drifts in units of $k_B T/ns$.

| Molecule: | barnase | barnase | barnase | barnase | barnase | ubiquitin | ubiquitin |
|---|---|---|---|---|---|---|---|
| Ensemble: | NVE | NVE | NPT | NPT | NPT | NPT | NPT |
| T (K) | - | - | 298 | 298 | 400 | 298 | 298 |
| Tolerance \Buffer | $5{\cdot}10^{-3}$ | $5{\cdot}10^{-5}$ | $5{\cdot}10^{-3}$ | $5{\cdot}10^{-5}$ | $5{\cdot}10^{-5}$ | $5{\cdot}10^{-3}$ | $5{\cdot}10^{-5}$ |
| $3.14{\cdot}10^{-4}$ | -0.536 | -0.601 | -0.353 | -0.750 | -0.937 | -0.277 | -0.828 |
| $10^{-4}$ | -0.125 | -0.236 | 0.313 | -0.074 | -0.203 | 0.416 | -0.036 |
| $3.14{\cdot}10^{-5}$ | 0.152 | -0.018 | 0.471 | 0.080 | 0.103 | 0.680 | 0.124 |
| $10^{-5}$ | 0.204 | 0.031 | 0.604 | 0.125 | 0.128 | 0.722 | 0.167 |
| $10^{-6}$ | 0.187 | 0.017 | 0.595 | 0.173 | 0.111 | 0.712 | 0.260 |
| $10^{-7}$ | 0.187 | 0.017 | 0.594 | 0.115 | 0.111 | 0.711 | 0.152 |
| $10^{-8}$ | 0.188 | 0.017 | 0.590 | 0.200 | 0.108 | 0.713 | 0.261 |
| $10^{-10}$ | 0.188 | 0.017 | 0.508 | 0.115 | 0.110 | 0.709 | 0.153 |

##### B. Temperature evolution over time

In Fig 2, the time evolution of temperature corresponding to an NVE simulation is presented. This figure can be compared with Fig 3 of the main article. Their corresponding simulations have been set up identically except that for the simulation shown here only the H-bonds are constrained (all-bonds are constrained in simulations reported in Fig 3 of the main article). The curves display a qualitative trend similar to the one displayed in the main article, namely, low accuracy in satisfying the constraints leads to unstable temperatures.

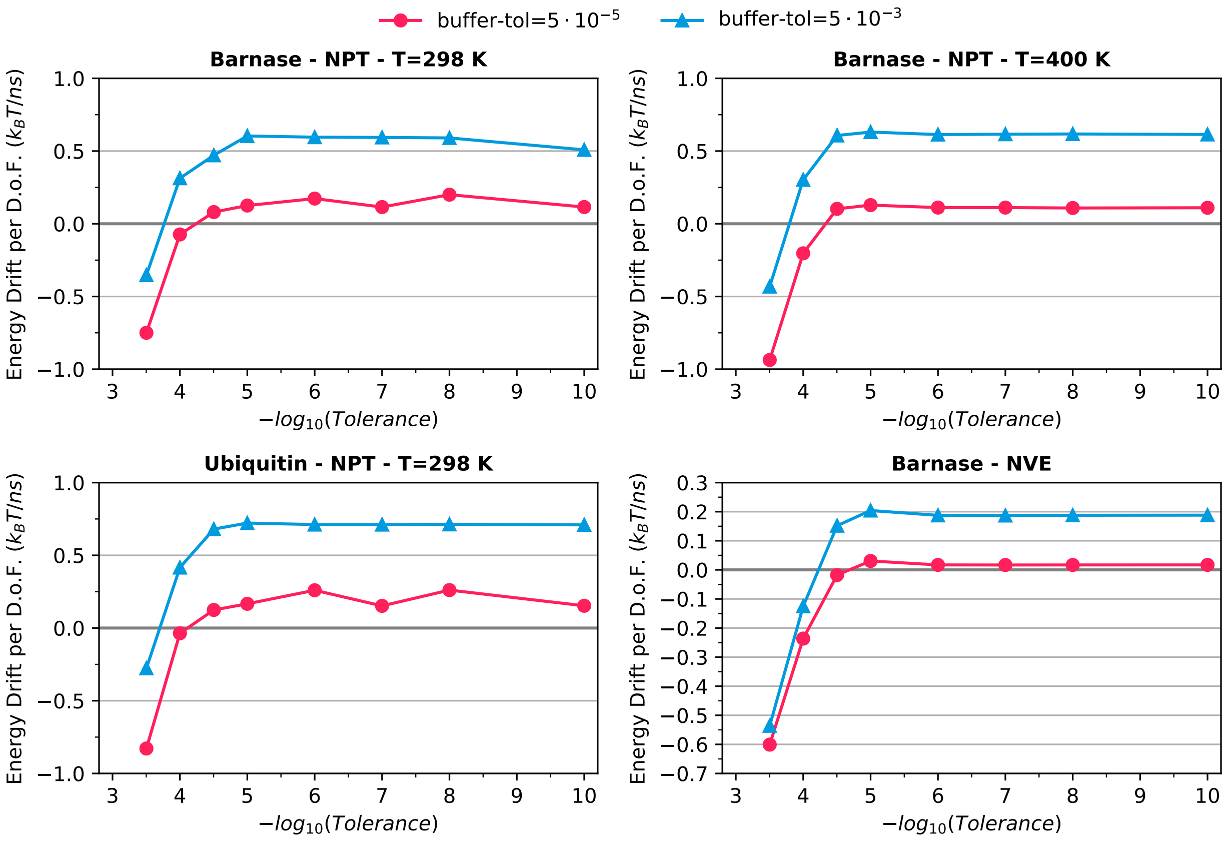


FIG. 1: Drifts of the conserved energy of the thermostat (NPT ensemble) and of the energy (NVE ensemble) per degree of freedom as a function of the tolerance in satisfying constraints imposed on H-bonds. Top: barnase in the NPT ensemble at T=298 K (left) and T=400 K (right); Bottom: ubiquitin in the NPT ensemble at T=298 K (left) and barnase in the NVE ensemble (right).

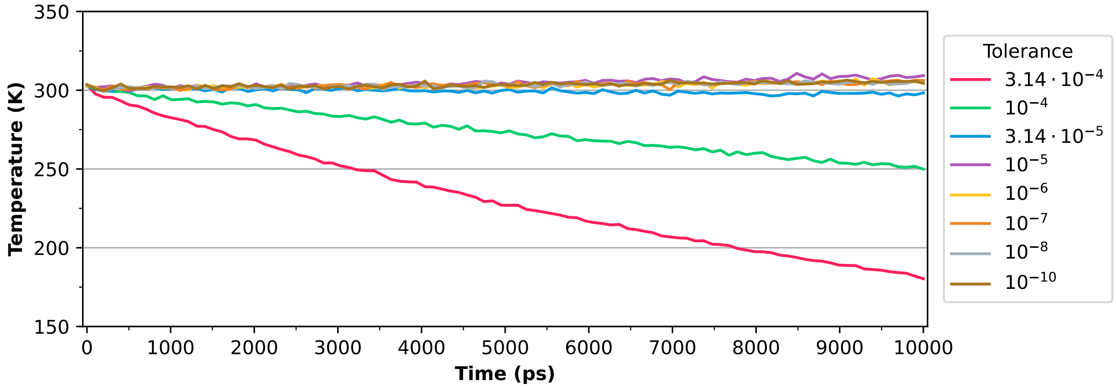


FIG. 2: Temperature evolution over time of barnase in the NVE ensemble using different tolerances in satisfying H-bonds constraints.

##### C. Distribution of errors in the constraints

In Fig 3, histograms of the time-averaged normalised errors of the constraints, defined as $\langle\, d_i \,\rangle := N_T^{-1} \sum_{k=1}^{N_T} d_i(t_k)$, are depicted. They correspond to simulations of ubiquitin (NPT ensemble, time step of 2 fs). The panel on the left corresponds to data obtained when setting the SHAKE algorithm, whereas the panel on the right corresponds to data obtained when setting P-LINCS. The rest of parameters in these simulations have been set to GROMACS default values. For both solvers most of the constraints errors have a positive average. The maximum values of $\langle\, d_i \,\rangle$ for the presented simulations are $7.4 \cdot 10^{-5}$ for SHAKE, and $1.1 \cdot 10^{-4}$ for P-LINCS.

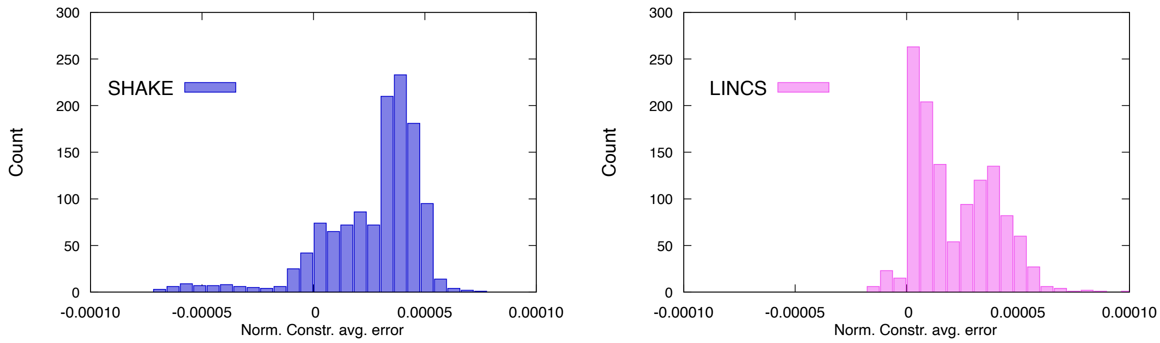


FIG. 3: Histograms of the time-averaged normalised errors of the constraints $\langle\, d_i \,\rangle$ for NPT simulations of ubiquitin using SHAKE (left) and P-LINCS (right).

In Figs. 4 and 5, normalised and non-normalised constraint errors obtained for one simulation of ubiquitin (NPT ensemble) with P-LINCS are depicted, respectively. The non-normalised constraint errors are defined with the same formulas as $\bar{d}$, $d_{Max}$ and $d_{min}$ (see the main article), though omitting $\sigma_i$ from the denominator of $d_i$. The P-LINCS algorithm was set with the default parameters in GROMACS (`lincs_order`=4, `lincs_iter`=2). The curves of normalised and non-normalised errors have a similar appearance, though their scales differ (and have different units). Analysis of Fig 4 and Fig 4 of the main article reveals that the constraint average error (blue lines) obtained when using P-LINCS is lower than that obtained with SHAKE. These averages are, however, positive (nonzero) for both solvers. Moreover, Fig 4 indicates that constraints with a violation (normalised error) above $10^{-4}$ can come out with P-LINCS, in contrast to what is observed with SHAKE. We do not display further examples of the errors of the lengths when using P-LINCS (apart from Figs 4 and 5) because for this solver the curves for all-bonds look qualitatively similar (smoother than SHAKE's, and barely having values well below $10^{-4}$).

Figs 6 to 13 display the normalised bond length error profiles over time of selected constrained bonds in a simulation of ubiquitin (SHAKE algorithm applied on all-bonds with a tolerance of $10^{-4}$, NPT ensemble at $298$ K and 1 atm). For the sake of comparisons, the error profiles are grouped by the type of bond. Selection includes bonds both from the main chain and the side chain of some amino acid residues. The analysis of these plots reveals that same types of bonds present similar error profiles, the overwhelming majority of which keeps positive over time for the conditions set in the simulations. Some bond length error plots also unveils time-correlated error profiles in similar constrained bonds when they appear in the same residue, as it is the case –for instance– of the five Carbon(CD1, CE1, CZ, CD2, CE2)-Hydrogen bonds of the aromatic ring of phenylalanines (Fig 11) or the case of the three Nitrogen(NZ)-Hydrogen bonds of the charged side chain amino group of lysines (compare panels `a-c` and `d-f` in Fig 12). Moreover, it can be observed that error profiles of some specific bonds, e.g. those formed between the side chain Oxygen atom (OG1) and its associated Hydrogen atom in residues of threonine (Fig 13), or the prior shown side chain Nitrogen(NZ)-Hydrogen bond in residues of lysine (Fig 12), present completely different shapes (lengthier regions of zero or quasi-zero error combined with smooth plateaus or valleys) to those shown for the majority of remaining bonds. The observed lower errors in these cases may indicate that bonds shared by one atom with a high charge (e.g. the polar atoms of side chains in some amino acid residues) and another one of opposite sign –like a Hydrogen– could be more averse to distortions of their bond distances because of the higher electrostatics implied in the interaction.

Figs. 6 to 13 display qualitatively different shapes for the errors. Many of them (e.g. 4, 5, 6) consist mainly of peaks. However, other are approximately constant (i.e. stay within a range of non-negligible values) for most of the time, like Fig 11-c, or Fig 12-f (low values). Others tend to switch between low values and maximum allowed values, like in panels `b`, `d` and `e` of Fig 11.

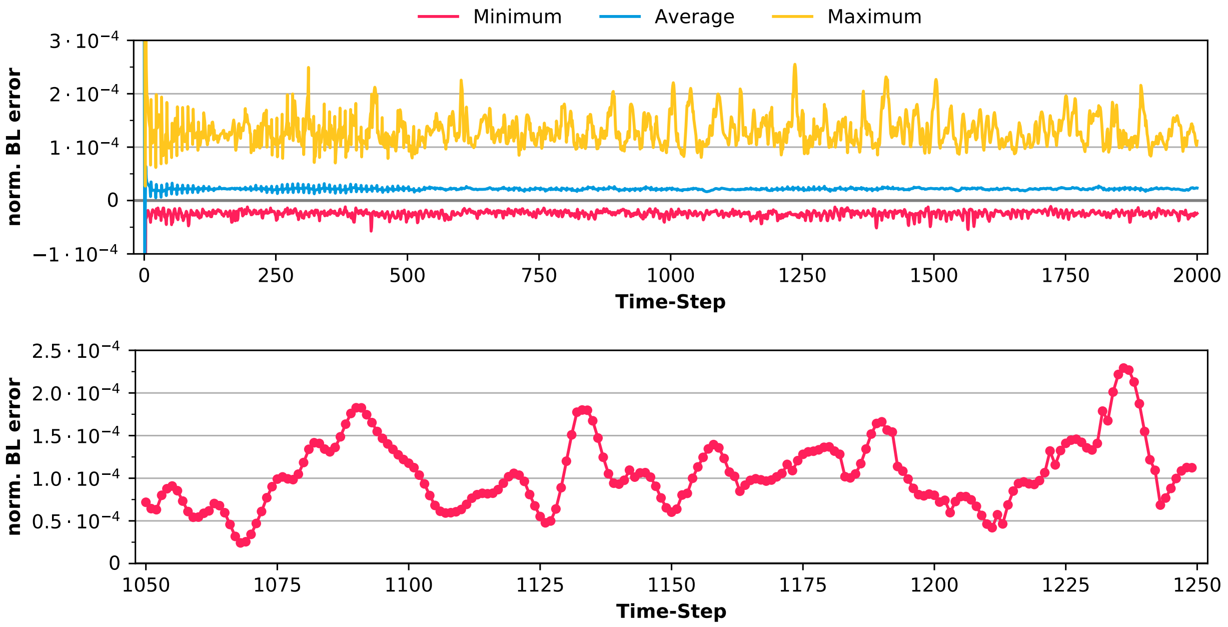


FIG. 4: Plots of the bond lengths normalised error over time in an MD run of ubiquitin with P-LINCS. The top panel includes the average, maximum and minimum values of the normalised error for all the constrained bonds (all-bonds). Truncated (not displayed) peaks reach 0.0020 and -0.0015. The bottom panel shows the normalised bond length error for a selected constrained bond (taken as an example).

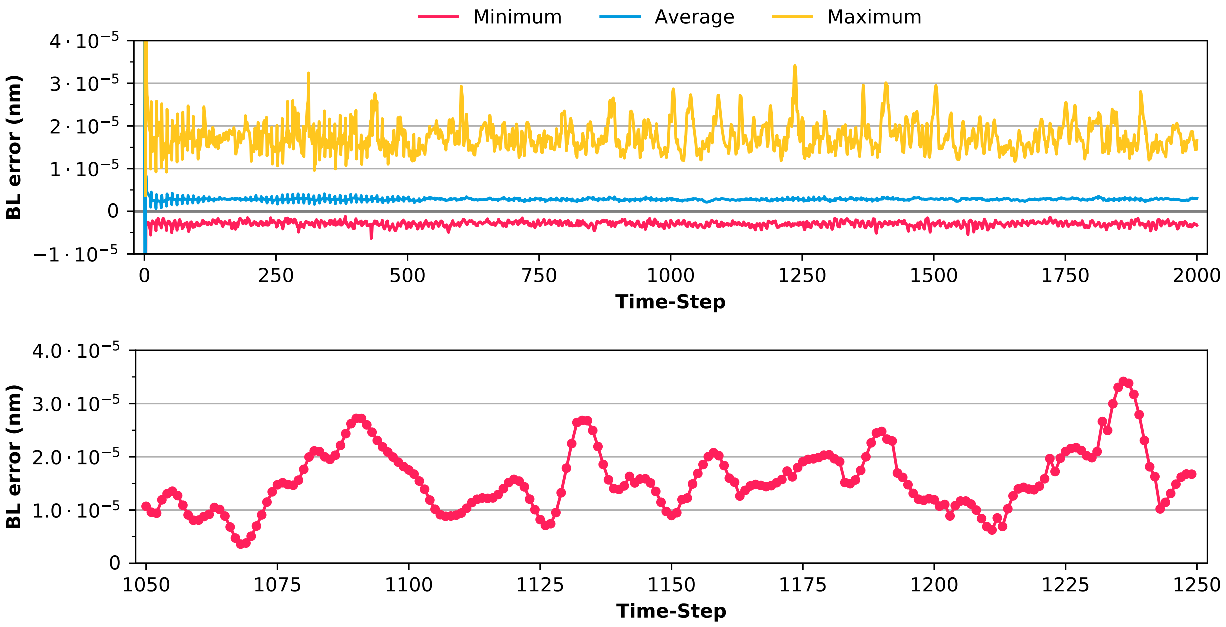


FIG. 5: Plots of the constraints error (difference between actual and expected bond lengths) over time for the same MD run of ubiquitin with P-LINCS used for Fig 5. The top panel includes the average, maximum and minimum values of the error for all the constrained bonds (all-bonds). Truncated (not displayed) peaks reach $2.7{\cdot}10^{-4}$ and $-2.1{\cdot}10^{-4}$. The bottom panel shows the bond length error for the selected constrained bond shown in Fig 5-bottom.

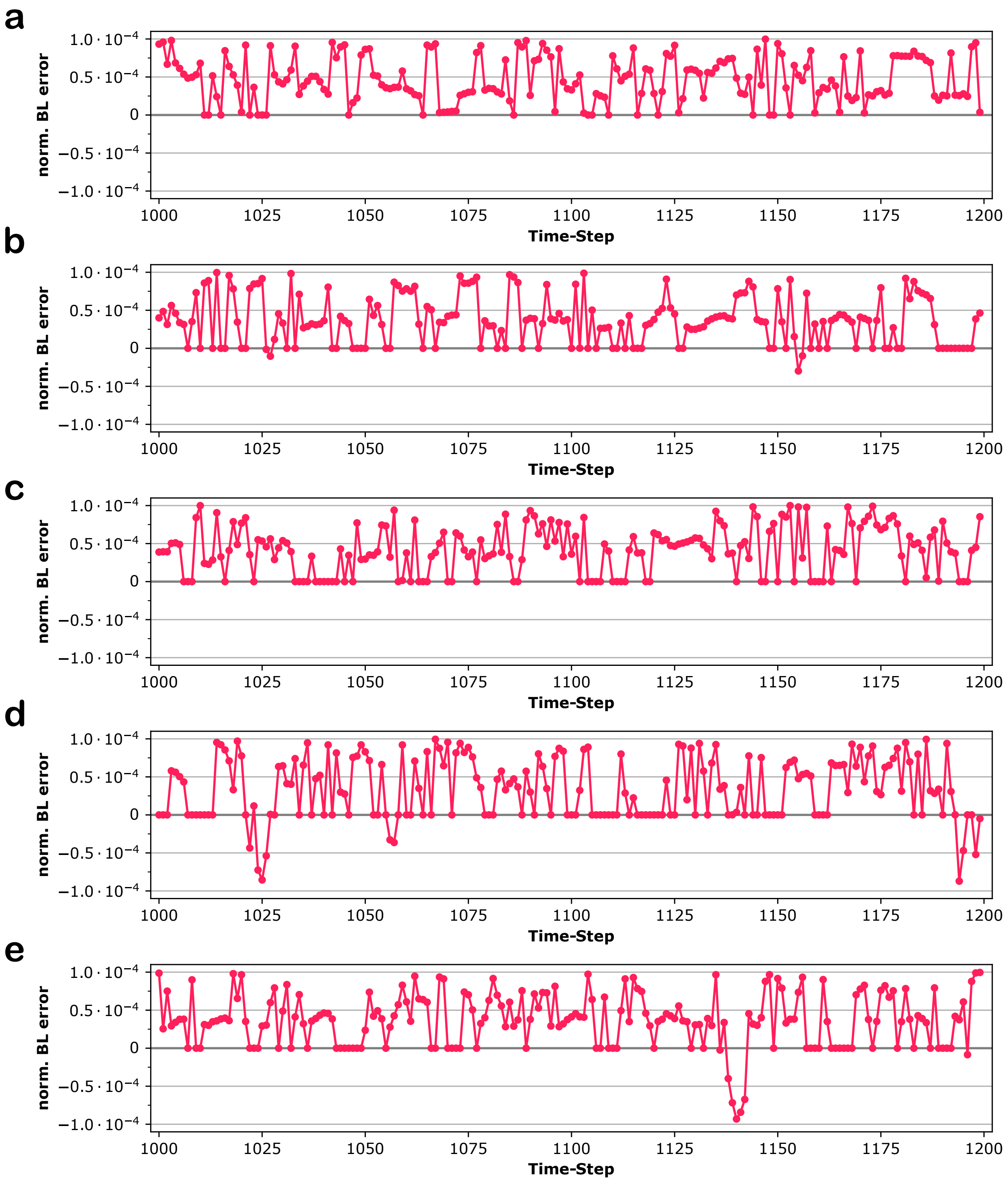


FIG. 6: Normalised bond length error as a function of time. Plots correspond to main chain Nitrogen(N)-Hydrogen bonds of the first five residues of ubiquitin in an NPT simulation with SHAKE algorithm (tolerance of $10^{-4}$, 298 K and 1 atm); a: Met1, b: Gln2, c: Ile3, d: Phe4, e: Val5.

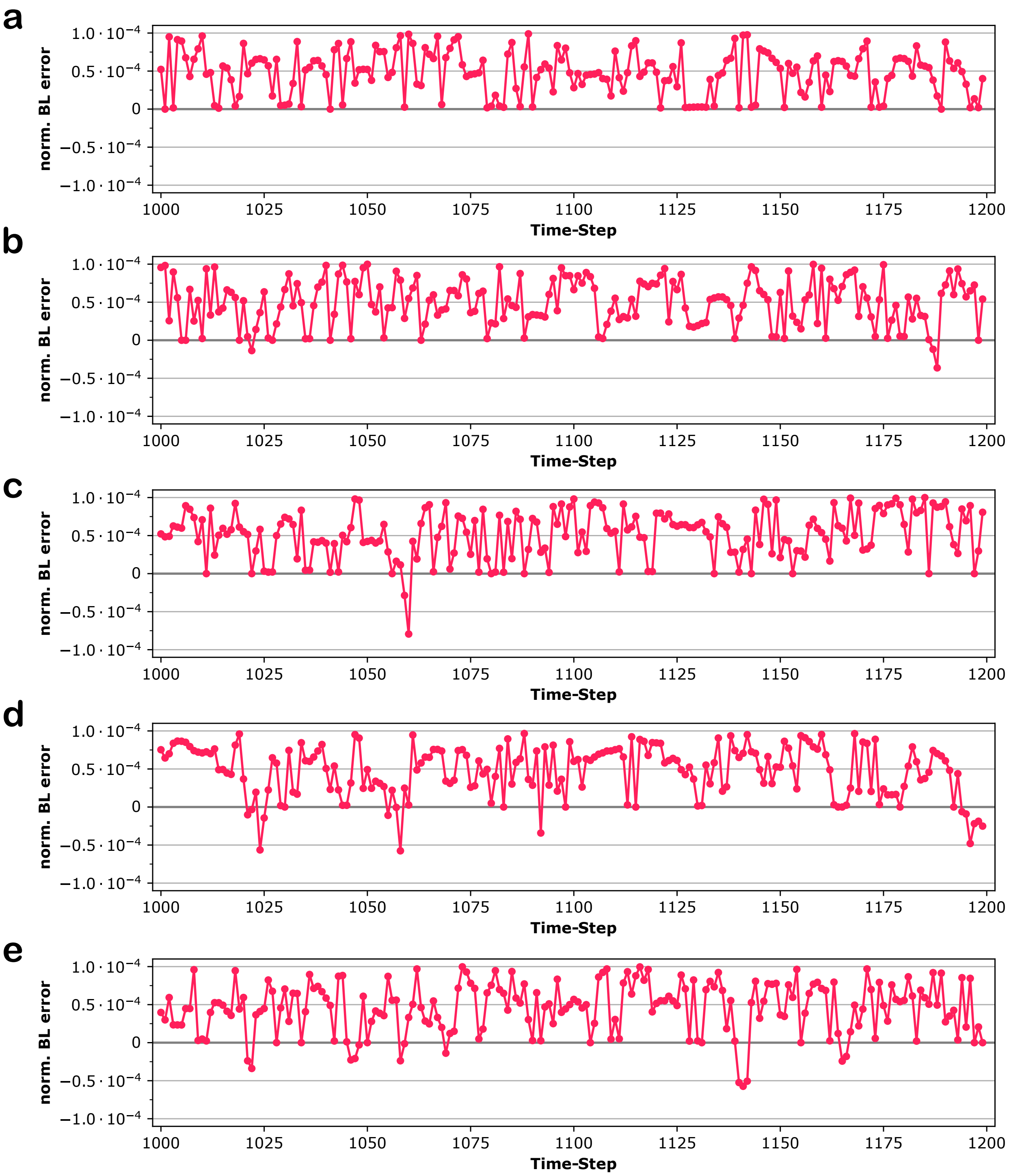


FIG. 7: Normalised bond length error as a function of time. Plots correspond to main chain Nitrogen(N)-Carbon(CA) bonds of the first five residues of ubiquitin in an NPT simulation with SHAKE algorithm (tolerance of $10^{-4}$, 298 K and 1 atm); a: Met1, b: Gln2, c: Ile3, d: Phe4, e: Val5.

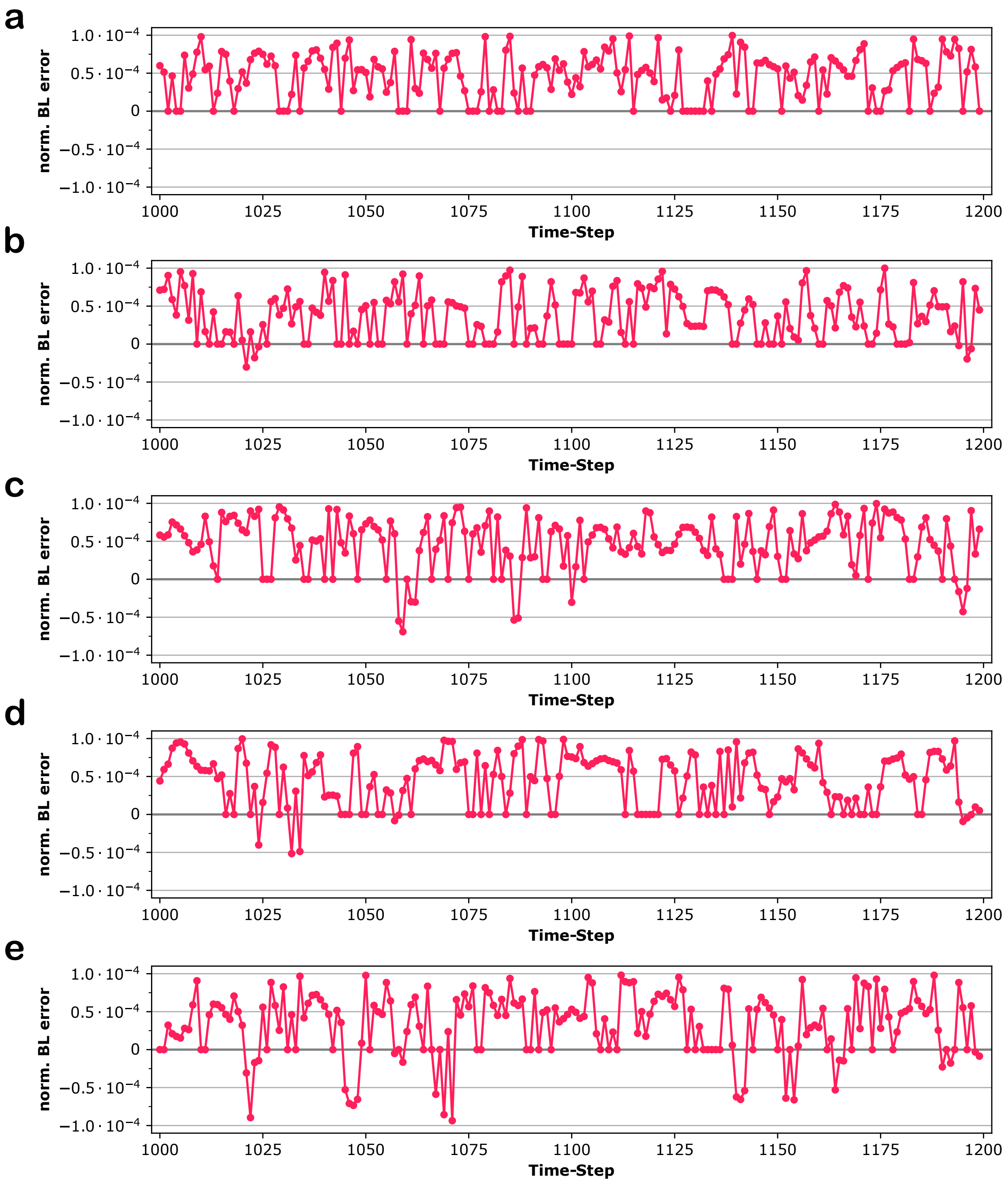


FIG. 8: Normalised bond length error as a function of time. Plots correspond to main chain Carbon(CA)-Hydrogen(HA) bonds of the first five residues of ubiquitin in an NPT simulation with SHAKE algorithm (tolerance of $10^{-4}$, 298 K and 1 atm); a: Met1, b: Gln2, c: Ile3, d: Phe4, e: Val5.

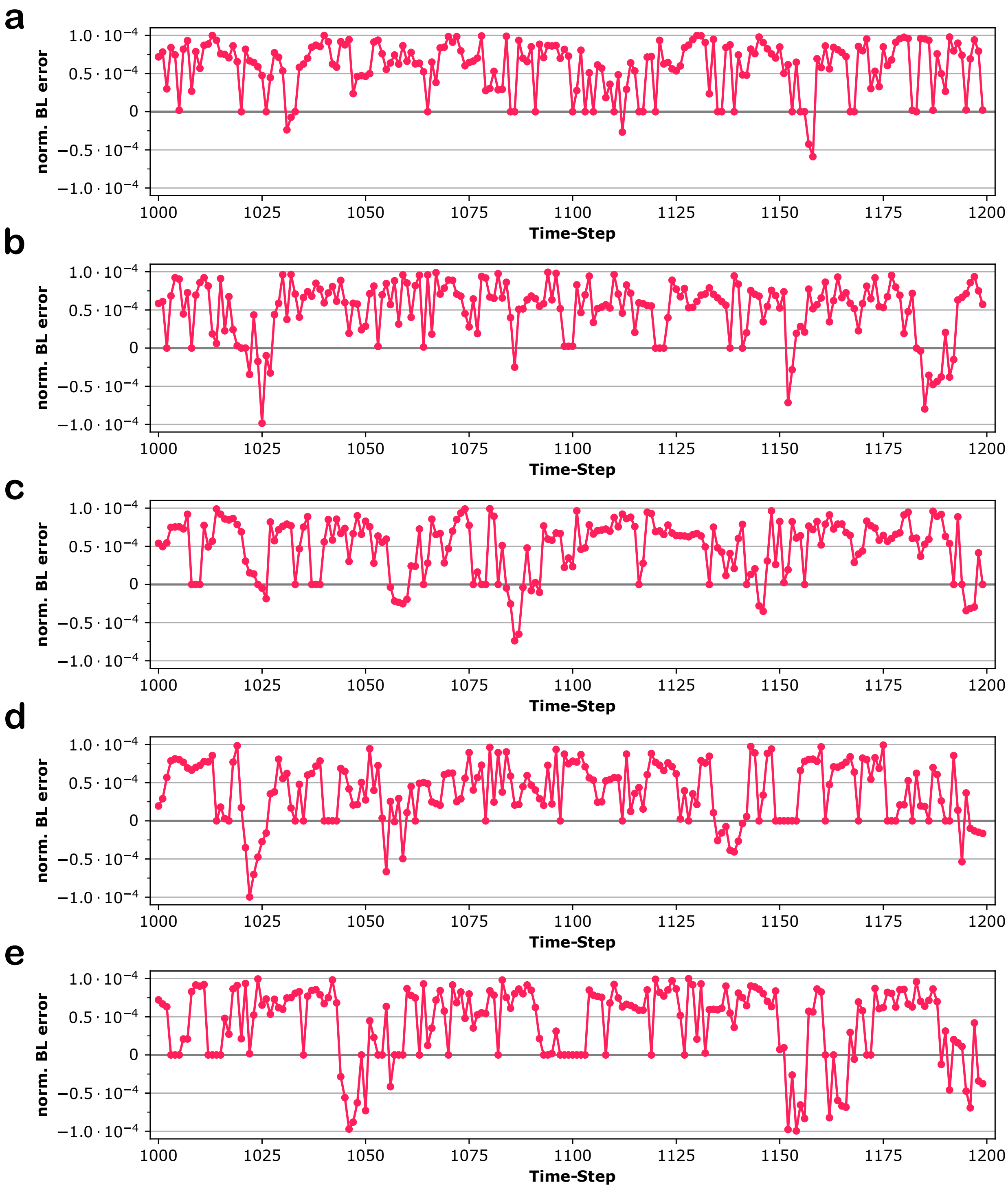


FIG. 9: Normalised bond length error as a function of time. Plots correspond to main chain Carbon(CA)-Carbon(C) bonds of the first five residues of ubiquitin in an NPT simulation with SHAKE algorithm (tolerance of $10^{-4}$, 298 K and 1 atm); a: Met1, b: Gln2, c: Ile3, d: Phe4, e: Val5.

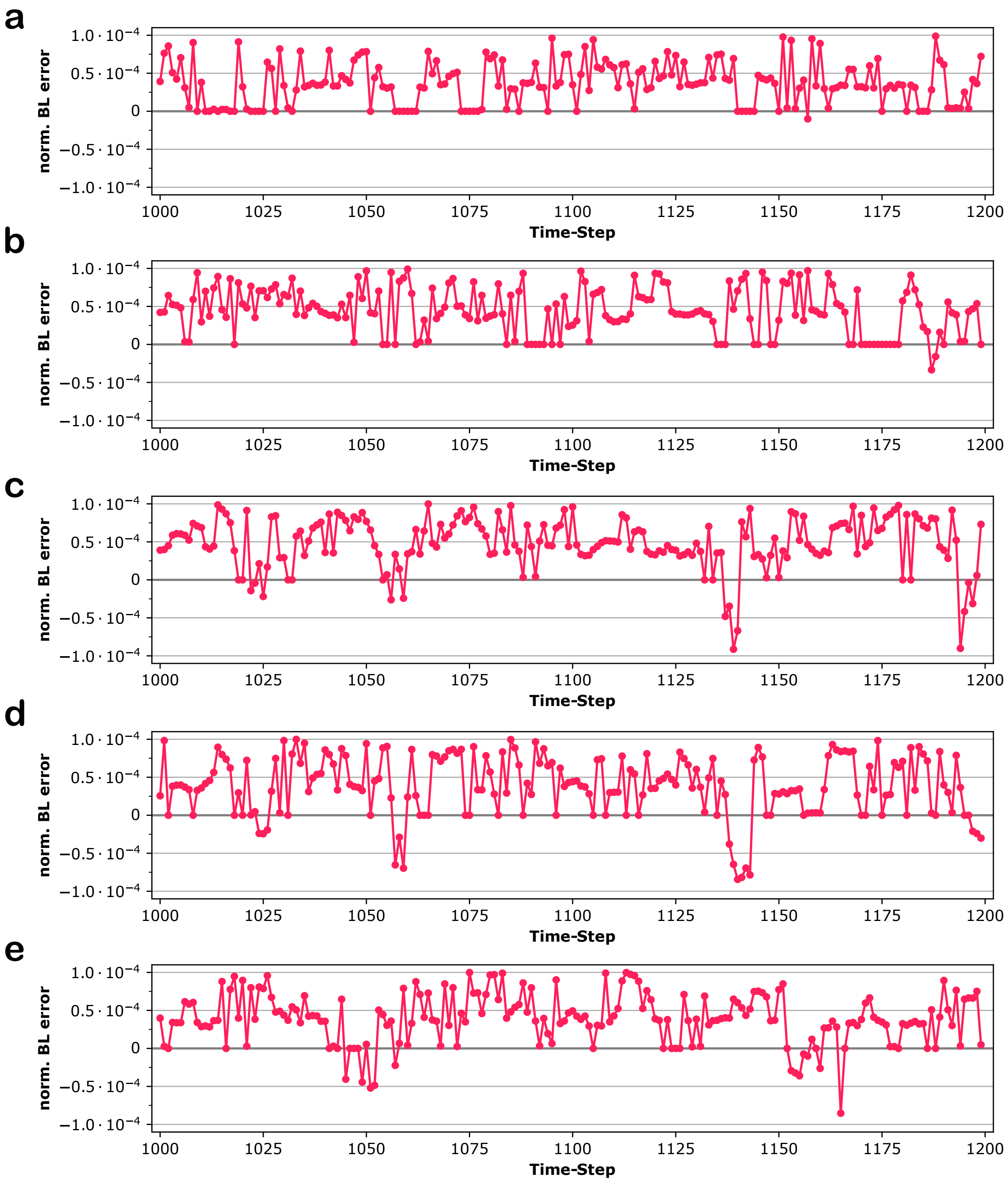


FIG. 10: Normalised bond length error as a function of time. Plots correspond to Carbon(C)-Nitrogen(N) peptidic bonds of the first five residues of ubiquitin in an NPT simulation with SHAKE algorithm (tolerance of $10^{-4}$, 298 K and 1 atm); a: Met1, b: Gln2, c: Ile3, d: Phe4, e: Val5.

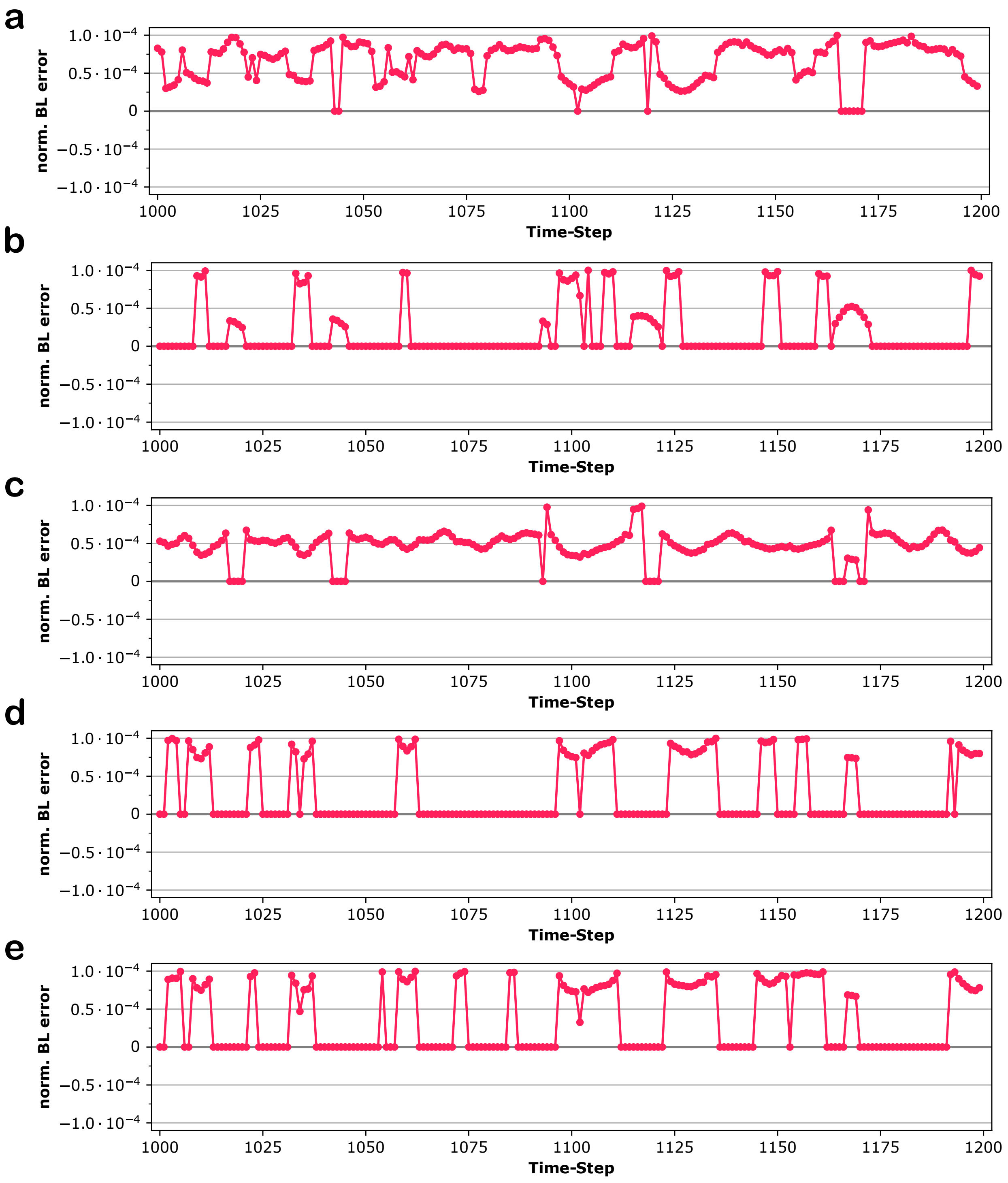


FIG. 11: Normalised bond length error as a function of time. Plots correspond to the five side chain Carbon-Hydrogen bonds of the aromatic ring of Phe4 in an NPT simulation of ubiquitin with SHAKE algorithm (tolerance of $10^{-4}$, 298 K and 1 atm); a: CD1-HD1, b: CE1-HE1, c: CZ-HZ, d: CD2-HD2, e: CE2-HE2.

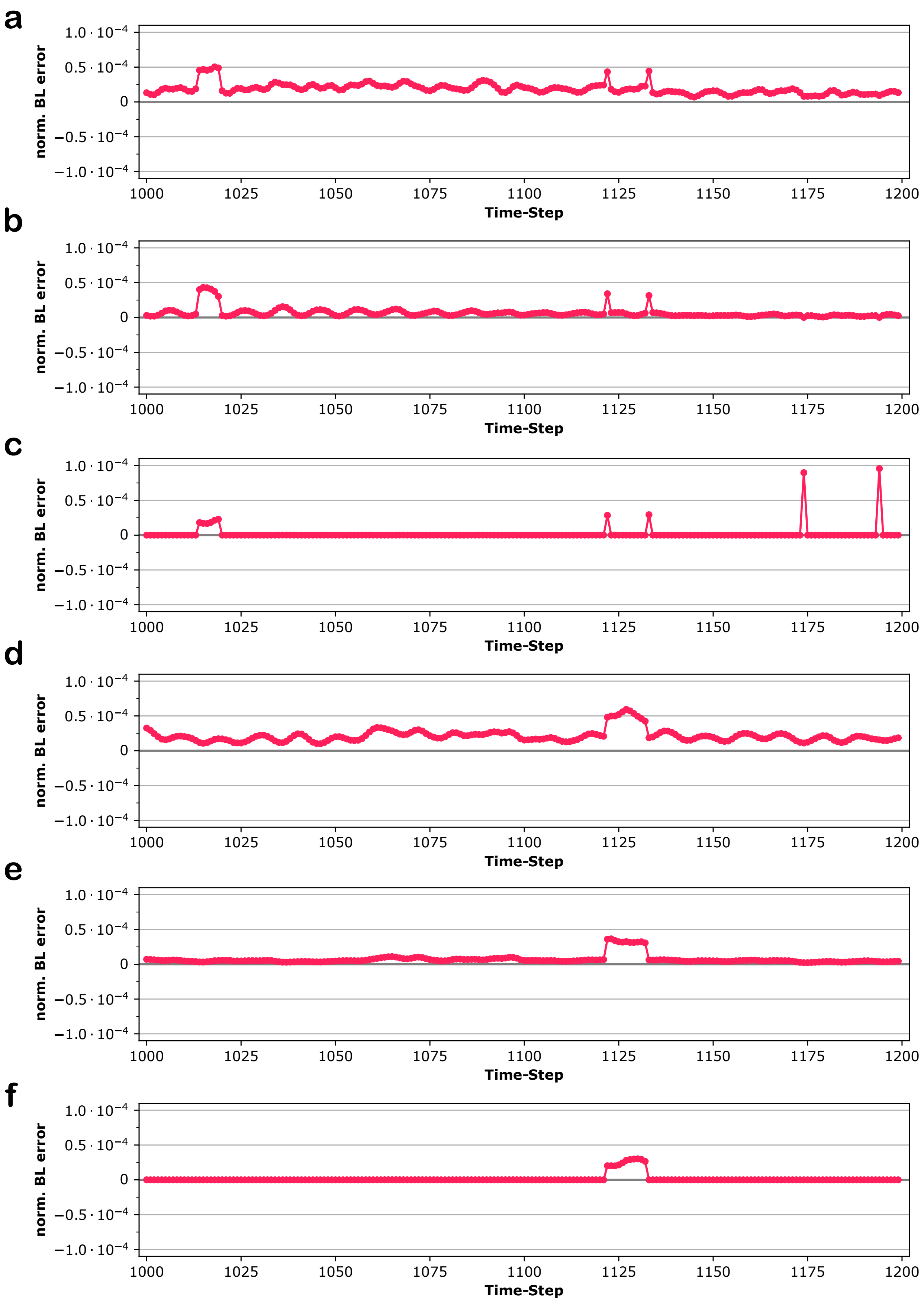


FIG. 12: Normalised bond length error as a function of time. Plots correspond to the three side chain Nitrogen(NZ)-Hydrogen bonds of the protonated amino group of two lysines in an NPT simulation of ubiquitin with SHAKE algorithm (tolerance of $10^{-4}$, 298 K and 1 atm); `a`: NZ-HZ1 in Lys6, `b`: NZ-HZ2 in Lys6, `c`: NZ-HZ3 in Lys6, `d`: NZ-HZ1 in Lys11, `e`: NZ-HZ2 in Lys11, `f`: NZ-HZ3 in Lys11.

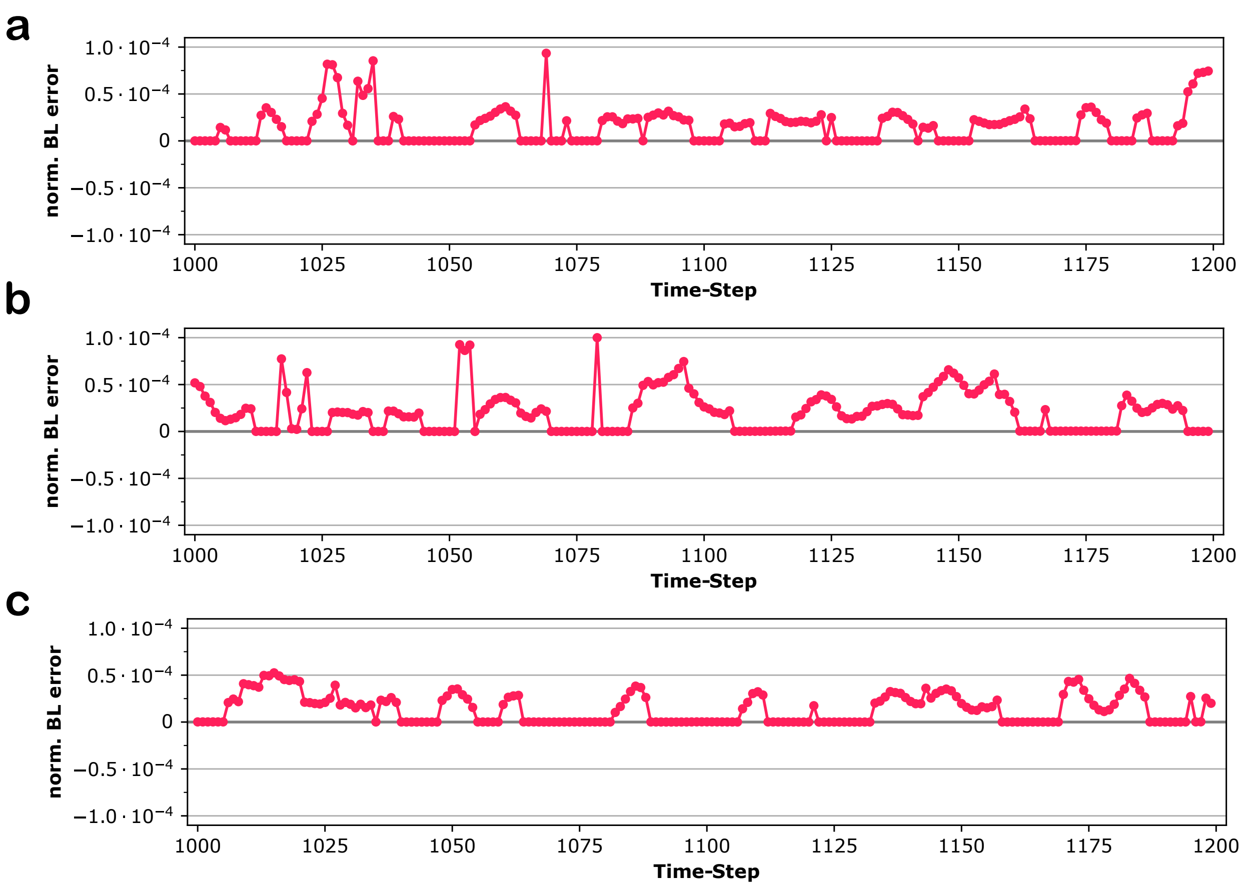


FIG. 13: Normalised bond length error as a function of time. Plots correspond to the side chain Oxygen(OG1)-Hydrogen bonds of three threonines in an NPT simulation of ubiquitin with SHAKE algorithm (tolerance of $10^{-4}$, 298 K and 1 atm); a: OG1-H in Thr7, b: OG1-H in Thr9, c: OG1-H in Thr12.

The autocorrelations of the degree of satisfaction of constraints (for consecutive time steps) are relatively high (see Fig 14). This can lead to inconsistencies given that for different stretches of the simulation the degree of satisfaction varies. The different degree of satisfaction of constraints in different simulation stretches implies different effective bond lengths throughout the simulations.

Since the energy of the molecule depends on the bond lengths [3], an inaccurate constraint solver can inject energy to the system, and doing so at a varying rate. Given that the total number of steps in a MD simulation is typically much higher than 1/tolerance when tolerance is set to 0.0001, one cannot guarantee that the errors in involved quantities will not accumulate till reaching unacceptable levels. This, however, would not happen if the tolerance were e.g. $10^{-13}$.

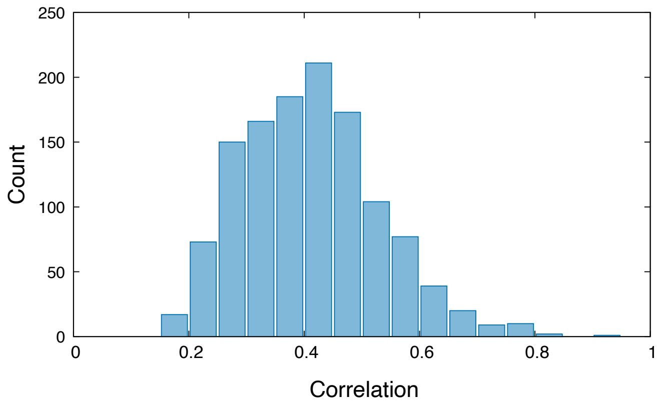


FIG. 14: Histogram of the autocorrelations (with the previous time step) of bond length errors for a selected constrained bond in a simulation of ubiquitin with constraints imposed on all atoms with SHAKE algorithm and a tolerance of $10^{-4}$ over 2000 time steps (NPT ensemble at 298 K and 1 atm).

#### II. RESULTS - PERFORMANCE

Tables III to VII summarise the results of our performance calculations for the three tested proteins (ubiquitin, COVID-19 main protease, human SSU-procesome) with different values of the constraint solver tolerance (`shake_tol`); *all* stands for the total execution time; *constr* corresponds to the time spent to solve the constraints (all bonds) on the protein (not the solvent). All timings are in seconds. The results presented in Tables III to VII indicate that ILVES-PC performs better than the other solvers for all the analyzed cases, except when executing with 1 and 6 threads at the largest tolerance ($10^{-4}$). For these exceptions, the excess time of ILVES-PC is at most 0.05% of the total execution time (i.e. negligible). Note that in contrast to what happens with SHAKE, the average error of the results obtained with ILVES and a tolerance of $10^{-4}$ is well below that threshold.

Fig 15 presents the results of the executions using 24 and 48 parallel threads in simulations where the cutoffs for the electrostatic and van der Waals interactions were set to 1.2 nm (in contrast to the results presented in the main text wherein these cutoffs were set to 1.0 nm).

TABLE III: ONE THREAD (serial execution): Total execution time and time required by the constraint solver for different accuracy limits (tolerances) and three different proteins.

| | Tolerance | $10^{-4}$ | | $10^{-6}$ | | $10^{-8}$ | | $10^{-10}$ | | $10^{-12}$ | |
|---|---|---|---|---|---|---|---|---|---|---|---|
| Protein | Solver | all | constr | all | constr | all | constr | all | constr | all | constr |
| UBIQ | SHAKE | 2080.7 | 16.5 | 2097.3 | 31.7 | 2096.6 | 47.4 | 2126.6 | 62.7 | 2136.7 | 78.7 |
| | P-LINCS-O4 | 2078.2 | 10.9 | 2090.6 | 21.2 | 2104.4 | 32.1 | 2102.2 | 42.2 | 2122 | 53 |
| | P-LINCS-O8 | 2060.1 | 10.9 | 2078.4 | 20.9 | 2075.3 | 31.5 | 2082.6 | 42.3 | 2120.6 | 52.8 |
| | ILVES-PC | 2075.5 | 11.7 | 2080 | 12.3 | 2073.4 | 16 | 2086.7 | 16.1 | 2079.7 | 16.2 |
| COVID | SHAKE | 7279.6 | 64.8 | 7282.5 | 117.8 | 7323.8 | 172.3 | 7364.6 | 227.1 | 7472.3 | 281.8 |
| | P-LINCS-O4 | 7200.6 | 43.5 | 7213.2 | 80.6 | 7260.8 | 119.4 | 7299.7 | 161.8 | 7407.1 | 202.7 |
| | P-LINCS-O8 | 7212.2 | 43.5 | 7262.2 | 80.6 | 7239.5 | 119.4 | 7361.6 | 161.6 | 7335.5 | 202.6 |
| | ILVES-PC | 7300.5 | 45.5 | 7199.1 | 61 | 7232.8 | 63 | 7242.2 | 62.9 | 7246.3 | 69.7 |
| SSU-PR | SHAKE | 27941.5 | 123.7 | 28224.4 | 219.8 | 28021.1 | 318.3 | 28222.1 | 417.4 | 28579.8 | 518.6 |
| | P-LINCS-O4 | 27755.9 | 79.4 | 27911.6 | 156.8 | 27946.9 | 242.3 | 28104.5 | 321.9 | 28634.5 | 406.4 |
| | P-LINCS-O8 | 27836.7 | 79.5 | 28041.2 | 156.8 | 27997.1 | 242.4 | 27966.3 | 321 | 28447.3 | 405.8 |
| | ILVES-PC | 27772.9 | 81.7 | 28150.2 | 117.9 | 27789.4 | 120.6 | 27859.1 | 112.2 | 28296.9 | 143.1 |


[1] M. Abraham, B. Berk Hess, D. van der Spoel, and E. Lindahl, *GROMACS user manual version 5.0.4* (http://www.gromacs.org/Documentation/Manual, 2014).

[2] P. Eastman and V. S. Pande, bioRxiv (2016), https://www.biorxiv.org/content/early/2016/10/24/083055.full.pdf, URL `https://www.biorxiv.org/content/early/2016/10/24/083055`.

[3] C. Oostenbrink, A. Villa, A. E. Mark, and W. F. van Gunsteren, J. Comput. Chem **26**, 1719 (2005).

TABLE IV: 6 THREADS: Total execution time and time required by the constraint solver for different accuracy limits (tolerances) and three different proteins.

| | Tolerance | $10^{-4}$ | | $10^{-6}$ | | $10^{-8}$ | | $10^{-10}$ | | $10^{-12}$ | |
|---|---|---|---|---|---|---|---|---|---|---|---|
| Protein | Solver | all | constr | all | constr | all | constr | all | constr | all | constr |
| UBIQ | SHAKE | 403.8 | 15.8 | 417.8 | 31.2 | 436.3 | 44.6 | 443.7 | 59.2 | 466.7 | 74.1 |
| | P-LINCS-O4 | 391.3 | 3.9 | 401.7 | 7.4 | 400.8 | 12 | 398.6 | 14.5 | 407.5 | 21.8 |
| | P-LINCS-O8 | 392.6 | 4.2 | 395 | 8 | 395.9 | 12 | 398.9 | 15 | 404.4 | 19.2 |
| | ILVES-PC | 391.4 | 4.4 | 392.1 | 4.4 | 384.3 | 5.7 | 393.2 | 5.7 | 389.5 | 5.4 |
| COVID | SHAKE | 1374.7 | 64.7 | 1423.2 | 117.8 | 1481.6 | 172.1 | 1540.3 | 226.7 | 1587.1 | 281.5 |
| | P-LINCS-O4 | 1317.1 | 9.8 | 1326.4 | 17.7 | 1332.2 | 25.8 | 1352.3 | 35.3 | 1350.1 | 43.7 |
| | P-LINCS-O8 | 1315.1 | 9.8 | 1325.2 | 17.9 | 1335.3 | 25.7 | 1340.6 | 34.7 | 1360.4 | 43.6 |
| | ILVES-PC | 1321.2 | 9.9 | 1327.3 | 13.4 | 1325.7 | 13.4 | 1317.7 | 13.3 | 1342.6 | 15.5 |
| SSU-PR | SHAKE | 5120.7 | 123.8 | 5199.6 | 219.8 | 5323.7 | 318.7 | 5416 | 418 | 5632.8 | 518.7 |
| | P-LINCS-O4 | 5008.2 | 14.5 | 5031.8 | 27.3 | 5048.3 | 42 | 5055.8 | 55 | 5053.5 | 68.5 |
| | P-LINCS-O8 | 5019 | 14.4 | 5031.2 | 27.5 | 5035.8 | 41.8 | 5042.3 | 54.7 | 5060.4 | 68.7 |
| | ILVES-PC | 5030.4 | 14.7 | 5102.6 | 19.8 | 5009.2 | 19.8 | 5015.8 | 19.8 | 5033.5 | 24.9 |

TABLE V: 12 THREADS: Total execution time and time required by the constraint solver for different accuracy limits (tolerances) and three different proteins.

| | Tolerance | $10^{-4}$ | | $10^{-6}$ | | $10^{-8}$ | | $10^{-10}$ | | $10^{-12}$ | |
|---|---|---|---|---|---|---|---|---|---|---|---|
| Protein | Solver | all | constr | all | constr | all | constr | all | constr | all | constr |
| UBIQ | SHAKE | 230.8 | 15.8 | 240.4 | 29.7 | 259.1 | 44.5 | 273.7 | 59 | 290.1 | 73.5 |
| | P-LINCS-O4 | 213.7 | 4.7 | 222.5 | 9.4 | 225 | 14.1 | 230.5 | 19.3 | 237.6 | 24.7 |
| | P-LINCS-O8 | 216.6 | 4.8 | 222.5 | 9.8 | 227.5 | 14.2 | 230.5 | 19.1 | 236.8 | 24 |
| | ILVES-PC | 218.4 | 4.6 | 216.8 | 4.5 | 215.6 | 5.6 | 217.3 | 5.9 | 216.9 | 6 |
| COVID | SHAKE | 762.4 | 64.6 | 822 | 117.6 | 867.5 | 172 | 929.2 | 226.8 | 976.8 | 281.4 |
| | P-LINCS-O4 | 705.6 | 8.1 | 706.6 | 14.8 | 718.5 | 22.1 | 723.4 | 30.1 | 726.6 | 36.8 |
| | P-LINCS-O8 | 700.4 | 8.1 | 708.2 | 14.9 | 716.3 | 21.8 | 721.7 | 29.6 | 723.7 | 36.3 |
| | ILVES-PC | 698.4 | 7.2 | 704.6 | 9.8 | 708.8 | 10.5 | 707.4 | 10.5 | 707.1 | 11.1 |
| SSU-PR | SHAKE | 2724.9 | 123.7 | 2821.2 | 219.9 | 2946.9 | 319 | 3044.2 | 418.1 | 3136.2 | 518.3 |
| | P-LINCS-O4 | 2659.9 | 8.6 | 2638.1 | 15.5 | 2634 | 22.6 | 2653.6 | 29.7 | 2671.6 | 37.2 |
| | P-LINCS-O8 | 2630.3 | 8.3 | 2619.3 | 14.9 | 2640 | 22.8 | 2630.9 | 29.6 | 2651.5 | 37.4 |
| | ILVES-PC | 2675.1 | 8.1 | 2631.7 | 10.6 | 2612.3 | 10.7 | 2626.5 | 10.7 | 2633.3 | 13.5 |

TABLE VI: 24 THREADS: Total execution time and time required by the constraint solver for different accuracy limits (tolerances) and three different proteins.

| | Tolerance | $10^{-4}$ | | $10^{-6}$ | | $10^{-8}$ | | $10^{-10}$ | | $10^{-12}$ | |
|---|---|---|---|---|---|---|---|---|---|---|---|
| Protein | Solver | all | constr | all | constr | all | constr | all | constr | all | constr |
| UBIQ | SHAKE | 136.8 | 15.8 | 149.7 | 29.8 | 168.6 | 44.5 | 185.1 | 59.2 | 200.5 | 73.9 |
| | P-LINCS-O4 | 130.5 | 5 | 131.2 | 8.9 | 139.5 | 14.4 | 144.1 | 19.4 | 145.3 | 22.8 |
| | P-LINCS-O8 | 131.1 | 5.1 | 129.3 | 8.8 | 139.2 | 14.2 | 140.2 | 18.4 | 148.4 | 23.3 |
| | ILVES-PC | 129.1 | 4.7 | 127.4 | 4.7 | 131.1 | 6.5 | 129.6 | 6.2 | 131.5 | 6.5 |
| COVID | SHAKE | 453 | 64.8 | 511.6 | 118.1 | 558.3 | 172.1 | 628.6 | 227 | 675.6 | 281.4 |
| | P-LINCS-O4 | 398.6 | 6.7 | 404.3 | 12 | 407.7 | 17.8 | 410.3 | 23 | 420.8 | 29.8 |
| | P-LINCS-O8 | 397.4 | 6.7 | 398.5 | 12 | 409.8 | 18 | 398.7 | 22 | 419.2 | 29.8 |
| | ILVES-PC | 399.4 | 6.4 | 405.4 | 8.2 | 393.4 | 7.8 | 398.6 | 8.3 | 401.1 | 10 |
| SSU-PR | SHAKE | 1569.1 | 123.9 | 1665.5 | 219.8 | 1788.9 | 319.1 | 1888.4 | 418.3 | 1999.3 | 518.8 |
| | P-LINCS-O4 | 1462.6 | 5.5 | 1458.3 | 9.4 | 1480.5 | 13.1 | 1458.3 | 17.4 | 1468.2 | 21.5 |
| | P-LINCS-O8 | 1440.3 | 5.3 | 1446.7 | 9 | 1460.3 | 13 | 1455.9 | 17.1 | 1512.5 | 21.6 |
| | ILVES-PC | 1448 | 5.1 | 1455.2 | 6.9 | 1480.9 | 7.2 | 1459.7 | 6.9 | 1465.3 | 8.5 |

TABLE VII: 48 THREADS: Total execution time and time required by the constraint solver for different accuracy limits (tolerances) and three different proteins.

| | Tolerance | $10^{-4}$ | | $10^{-6}$ | | $10^{-8}$ | | $10^{-10}$ | | $10^{-12}$ | |
|---|---|---|---|---|---|---|---|---|---|---|---|
| Protein | Solver | all | constr | all | constr | all | constr | all | constr | all | constr |
| UBIQ | SHAKE | 96.1 | 16.6 | 107.3 | 31.3 | 124.7 | 46.9 | 140.6 | 61.5 | 157.6 | 76.4 |
| | P-LINCS-O4 | 91.3 | 13.2 | 105.8 | 27.2 | 96.2 | 20.5 | 102.5 | 27.6 | 122.4 | 45.4 |
| | P-LINCS-O8 | 93.1 | 13.3 | 108.2 | 26.9 | 95.9 | 20.2 | 102.7 | 27.7 | 111.3 | 33.3 |
| | ILVES-PC | 87.2 | 8.6 | 81.7 | 6 | 92.3 | 12.9 | 91.6 | 11.9 | 86.3 | 10.9 |
| COVID | SHAKE | 291 | 67.2 | 348.9 | 120.4 | 401.8 | 174.8 | 465.2 | 229.6 | 510.8 | 284.5 |
| | P-LINCS-O4 | 244.5 | 14.5 | 256.7 | 28.6 | 270.5 | 44.9 | 250.8 | 30 | 303.9 | 76.5 |
| | P-LINCS-O8 | 245.3 | 14.6 | 251.5 | 27.9 | 273.6 | 44.5 | 257.9 | 31 | 273.2 | 39.2 |
| | ILVES-PC | 236.9 | 10.8 | 231 | 9.3 | 236.8 | 14.7 | 237.6 | 15.5 | 241.4 | 11 |
| SSU-PR | SHAKE | 995.4 | 124.5 | 1088.4 | 221.4 | 1196.7 | 319.8 | 1291.4 | 419 | 1393.4 | 520 |
| | P-LINCS-O4 | 874.1 | 4.2 | 875.5 | 7 | 896.2 | 15 | 886.1 | 19.6 | 884.3 | 16.8 |
| | P-LINCS-O8 | 874.3 | 5.2 | 880.7 | 10.3 | 903.5 | 15.1 | 886.2 | 13.3 | 894.5 | 16.5 |
| | ILVES-PC | 870.3 | 3.7 | 874.6 | 5.3 | 887.8 | 6.2 | 890.5 | 5 | 875.1 | 6.8 |

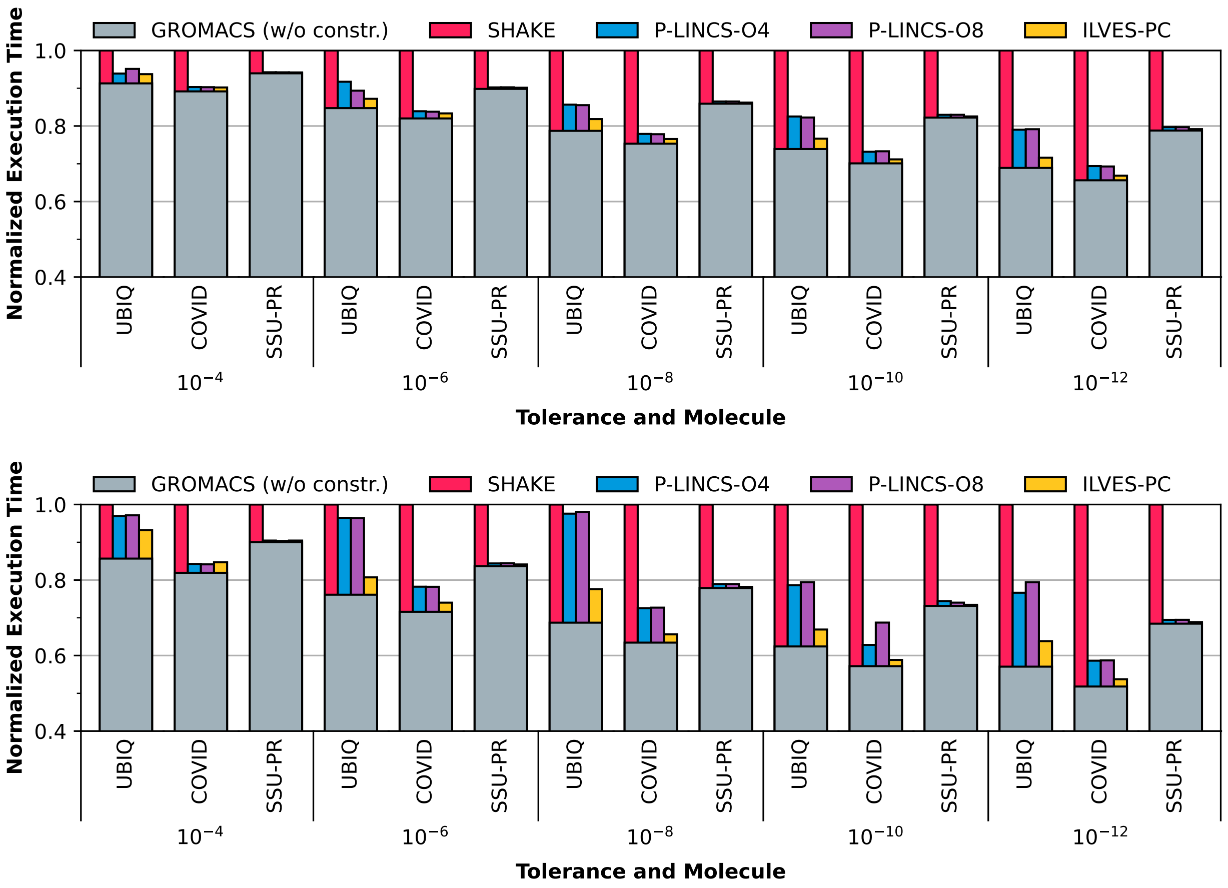


FIG. 15: Normalized execution times for different accuracy limits (tolerances) and three different proteins. The height of the thick grey bars represents the execution time of GROMACS excluding the constraints computations. The height of the magenta, blue, purple, and yellow bars represent the time required by the different constraint solvers. Top: parallel execution with 24 threads; bottom: ibid. with 48 threads. Electrostatic cutoff of 1.2 nm. Note that the $y$-axis starts at $y = 0.4$ rather than $y = 0$. This has been done to emphasize the differences between the constraint solvers.
